# Stimulus category is associated with conscious access during motion-induced blindness

**DOI:** 10.64898/2026.10.16.759106

**Authors:** András Sárközy, Sophia Gommlich, Jonathan E. Robinson, Gyula Kovács

## Abstract

Stimuli from different categories, such as faces or letters, differ in semantic content and neural representation. How stimulus category relates to conscious access during perceptual suppression has been studied mainly with binocular rivalry and continuous flash suppression, in which suppression arises from interocular competition. Motion-induced blindness (MIB) is a form of perceptual suppression without interocular competition, in which a static peripheral target intermittently disappears from visual awareness in the presence of a continuously moving background. Here, we tested whether target category affects the temporal dynamics of MIB using human faces, familiar Roman letters, and phase-randomized noise images as targets. Faces disappeared more frequently than noise and showed shorter invisible and visible episodes, whereas overall invisibility fraction and latency to first disappearance did not differ reliably. Letters showed a smaller increase in disappearance rate relative to noise, with no reliable difference in invisible episode duration. These findings reveal systematic category-related differences in the temporal organization of MIB and identify associations between stimulus category and the dynamics of conscious access.

## Introduction

Motion-induced blindness (MIB) is a well-known visual illusion in which high-contrast peripheral target stimuli intermittently disappear from conscious perception when a moving background is superimposed on them (Bonneh et al., 2001). The perceptual suppression of stationary targets against the motion background has been used as a model for studying the neural dynamics of visual conscious access and awareness (Dieter et al., 2015). Beyond its psychophysical merit, MIB is widely used in paradigms testing consciousness as it produces reliable fluctuations in subjective visibility, despite a physically constant target, showing a clean dissociation between physical inputs and perceptual experiences (Donner et al., 2008; Schölvinck & Rees, 2010). In this respect, MIB complements other bistable stimuli and rivalry paradigms that are often exploited to isolate neural processes, specifically associated with conscious access rather than stimulus-driven bottom-up encoding mechanisms (Wang et al., 2013), a distinction also emphasized in predictive coding accounts of consciousness research (Hohwy & Seth, 2020; Seth & Hohwy, 2021).

Neuroimaging work further supports this view: subjective target disappearances during MIB are correlated with systematic changes in visual cortical activity that correspond to visibility, encompassing signals related to both the target and the mask within ventral and dorsal visual pathways (Donner et al., 2008), as well as robust responses that vary with visibility in retinotopically organized early visual areas (Schölvinck & Rees, 2010). Furthermore, MIB has also been used to study the extent to which visual processing and object updating can proceed without awareness, showing that changes to perceptually suppressed objects can still influence subsequent perceptual outcomes (Mitroff & Scholl, 2005). Finally, MIB is currently also used in a parallel experiment of our laboratories for testing the predictive-processing-based active inference theory of consciousness against other competing theories (Robinson et al., 2025), in the framework of a large adversarial collaboration network (INTREPID; https://www.templetonworldcharity.org/blog/integrated-information-theory-iit-and-predictive-processing-accounts-of-consciousness).

A significant body of research has identified several factors that modulate the strength and temporal properties of MIB in the past. These include low-level visual parameters such as target stimulus size, contrast, eccentricity, or spatial frequency (Bonneh et al., 2001; Hsu et al., 2004; Hsu et al., 2006; Bonneh et al., 2014; Wallis & Arnold, 2008). Target eccentricity modulates MIB disappearance dynamics, with more peripheral targets typically showing stronger perceptual suppression (Bonneh et al., 2001). This pattern is broadly consistent with cortical magnification values, since peripheral targets are represented with lower effective spatial resolution, which may increase susceptibility to disappearance under competing motion signals (Virsu & Rovamo, 1979). Our previous study also demonstrated anisotropies in the magnitude of MIB across various regions of the visual field, indicating that spatial stimulus location affects the stability of perceptual representations (Sárközy et al., 2024). In addition to these low-level physical stimulus properties, top-down factors such as prediction and attentional states are also key modulators of MIB dynamics (Chang et al., 2015; Schölvinck & Rees, 2009; Geng et al., 2007). Paradoxically, directing spatial attention towards the peripheral targets may increase the probability of their disappearance during MIB, suggesting stronger neural competition with the surrounding motion pattern and leading to more difficult conscious stimulus access (Schölvinck & Rees, 2009). In contrast, under conditions of reduced attention towards the target stimulus, perceptual transitions become less frequent, and periods where the stimuli are invisible are prolonged (Schölvinck & Rees, 2009; Geng et al., 2007).

Most previous studies of MIB have used targets with little object structure, typically small, high-contrast uniform dots or simple geometric forms, presented against a moving mask (Bonneh et al., 2001; Graf et al., 2002; Hsu et al., 2004; Hsu et al., 2006; Wallis & Arnold, 2008; Bonneh et al., 2014). This approach has been useful in identifying general factors that influence the illusory stimulus disappearance. However, it leaves open a related question that has received less attention, namely whether the category of the target affects the temporal dynamics of perceptual suppression or not. The question is relevant to MIB as a paradigm, since the illusion is widely used as a read-out of conscious access, including in experiments that compare theories of consciousness (Robinson et al., 2025). If its temporal dynamics depend on the identity of the suppressed target, then target category becomes a variable that such designs must control.

Human faces and letters of the known alphabet provide a useful comparison for examining this issue for several reasons. First, they both engage specific and well-characterized category-selective systems in the human brain (Kanwisher et al., 1997; Dehaene et al., 2002). Second, faces and letters differ in biological relevance and acquisition history: face processing builds on an early-emerging perceptual bias that is already measurable in newborns (Farroni et al., 2005; Frank et al., 2009), whereas letters are culturally acquired symbols that require explicit instruction and reshape ventral visual cortex during literacy acquisition (Dehaene & Cohen, 2007; Dehaene-Lambertz et al., 2018), supported by extensive visual expertise from our reading experiences (Dehaene et al., 2002; McCandliss et al., 2003). Finally, both categories can be discriminated at the small peripheral size used here, as tested under the static, no-inducer conditions of Experiment A, and are available in large sets whose low-level image statistics can be quantitatively characterized across exemplars. Behavioural studies further suggest that faces enjoy priority in selection and attention, as shown by efficient detection in visual search (Hershler & Hochstein, 2005) and strong effects on attentional engagement and disengagement (Bindemann et al., 2005; Palermo & Rhodes, 2007).

Neuroimaging studies have identified a left ventral occipitotemporal region that responds reliably to letter strings and words, with graded tuning consistent with hierarchical coding of orthographic structures (Dehaene et al., 2002; Vinckier et al., 2007). Electrophysiological evidence further indicates structured and sequential processing of letters and words in fusiform cortex (Thesen et al., 2012). At the behavioural level, letter identification is subject to well-described limits, related to feature efficiency and stimulus complexity (Pelli et al., 2006).

Whether the content of a suppressed stimulus shapes its own suppression has been examined extensively in binocular rivalry and continuous flash suppression (CFS). During rivalry, category-selective cortical responses follow the perceptual state of the observer rather than the physical input (Tong et al., 1998; Tong, Meng, & Blake, 2006), while no-report paradigms indicate that face-selective neurons encode both consciously perceived and suppressed stimuli (Hesse & Tsao, 2020). Under CFS, the face-inversion effect was larger for breaking suppression than for reverse-breaking suppression (Ciorli et al., 2025). In these paradigms, however, suppression arises from interocular competition. Whether stimulus category similarly shapes conscious access when suppression is induced without interocular competition, by a moving mask, has remained largely unexamined, so that MIB offers a mechanistically distinct test of the same question.

Current accounts of MIB have largely explained the perceptual suppression of the targets in terms of low-level sensory and decisional mechanisms. In line with this, MIB has been linked to changes in sensory evidence and shifts in decision criteria (Caetta et al., 2007), and has also been described as a scotoma-like discounting of stable targets embedded in global motion (New & Scholl, 2008). These accounts share a common form. Each is specified over the physical description of the display, and none makes an explicit category prediction once size and position are matched and measured intensity and contrast are closely comparable. The present study asks whether the remaining differences in image structure and category-related processing are associated with the temporal dynamics of MIB. Framed at this level, the question is not whether “faces are special”, but whether the representations that compete during MIB are adequately described in terms of image features alone, or whether the level at which a target category is encoded also contributes. The question is a property of the accounts rather than of any particular stimulus class, so the same question arises regardless of the stimulus categories actually used. At the same time, target disappearance during MIB is unlikely to reflect a complete loss of object representation, as previous work suggests that even suppressed objects can continue to be represented and updated outside awareness, and that changes introduced to the suppressed objects influence subsequent perceptual outcomes (Mitroff & Scholl, 2005). Related electrophysiological work suggests that neural activity related to stimulus category processing may remain detectable during MIB, consistent with the idea that stimulus reappearances reflect the change in perceptual access to an ongoing representation rather than a complete reinitiation of stimulus category processing (Harris et al., 2013).

MIB can vary along several partly dissociable temporal dimensions, including disappearance frequency, invisible-state or visible-state duration, cumulative invisibility, and latency to the first disappearance (Leopold & Logothetis, 1999; Brascamp et al., 2005). We therefore characterized the pattern across these measures rather than relying on disappearance rates alone. Joint changes in state durations and cumulative invisibility distinguish a change in the overall balance of suppression from a change in the tempo of perceptual alternation, whereas the latency of the first disappearance indexes the initial transition into suppression. The joint pattern is also informative about the physical description of the targets. A simple monotonic increase in suppression strength would be expected to shift the balance between the visible and invisible states and therefore cumulative invisibility. A pattern in which the state durations change while cumulative invisibility does not is not well described as a monotonic shift in suppression strength. It is more naturally described as a change in the tempo of alternation, in which the transition dynamics are altered in both directions while the balance between the two states is preserved.

The present study therefore examined whether MIB dynamics differ between faces, letters, and Fourier phase-randomized noise stimuli. We report two behavioural experiments. Experiment A served as a stimulus check and assessed whether the categories were discriminable at the target size and base peripheral location used in Experiment B, in the absence of the rotating inducer. Experiment B then used these stimulus categories in an MIB paradigm adapted from Sárközy et al. (2024) to assess whether target category influences the temporal dynamics of perceptual suppression. As current accounts of MIB do not make strong predictions specific to stimulus category, we expected overall invisibility fraction to vary little across conditions. Our prediction concerned the frequency of disappearances: based on evidence showing that category-related stimulus processing may persist during MIB (Harris et al., 2013), and that attentional factors influence disappearance dynamics (Schölvinck & Rees, 2009), we expected faces to show the highest disappearance rates, noise stimuli the lowest, and letters an intermediate rate. Any category effects on invisible episode duration were expected to be small. Visible episode duration was examined as an exploratory outcome, in order to characterize the temporal organization of the visible state.

## Methods and Materials

### Participants

A total of 39 participants took part in the present study. Experiment A included 5 female participants (mean age: 21.0 years, SD = 1.22, range 20–23 years). Experiment B included 34 participants (29 females, 5 males; mean age: 22.15 years, SD = 3.27, range 19–32 years). Recruitment was conducted through the internal system of the Institute of Psychology, Friedrich Schiller University Jena. All participants had normal or corrected-to-normal vision and reported no neurological or psychiatric history. Participation was rewarded with partial course credit.

Experiment A was designed as a stimulus discrimination check rather than as a formal hypothesis-testing experiment. Therefore, no separate a priori power analysis was conducted for Experiment A; instead, five participants completed repeated category discrimination trials, which indicated that the stimulus categories were discriminable at the target size and base peripheral location used in Experiment B, in the absence of the rotating inducer. For Experiment B, sample size was planned a priori using G*Power 3.1, based on the initially planned repeated measures analysis of the within-subject comparison across three stimulus categories. Assuming a medium effect size of f = 0.25, α = .05, power = .80, a correlation of .50 among repeated measures, and nonsphericity correction ε = 1, the analysis indicated a required sample size of 28 participants. The final sample of Experiment B included 34 participants, exceeding this planned requirement.

### Ethics

Ethical approval for the study was granted by the local ethics committee (ethics number: FSV 21/073), and all procedures were carried out in accordance with the Declaration of Helsinki. Written informed consent was obtained from each participant prior to testing.

### Paradigm

#### Experiment A

Each trial began with a central white fixation cross (500 ms) on a uniform grey background, followed by a peripheral target stimulus in the upper-left visual field at a centre eccentricity of 7.2 deg (583 pixels; Fig. 1b). The stimulus was displayed for 3 s. Participants pressed “x” for faces, “m” for letters, and withheld their response for noise images. A go/no-go mapping was used in order to limit response-selection demands, which increase with the number of stimulus-response alternatives (Hick, 1952; Hyman, 1953; Gomez et al., 2007; Proctor & Schneider, 2018). An inter-trial interval of 1 s separated successive trials. The experiment comprised three blocks of 45 trials each (135 trials total), with categories balanced and randomized within each block (15 faces, 15 letters, 15 noise images). Blocks were separated by 20 s rest periods with a centrally presented countdown, after which participants continued with a key press.

**Fig. 1.**
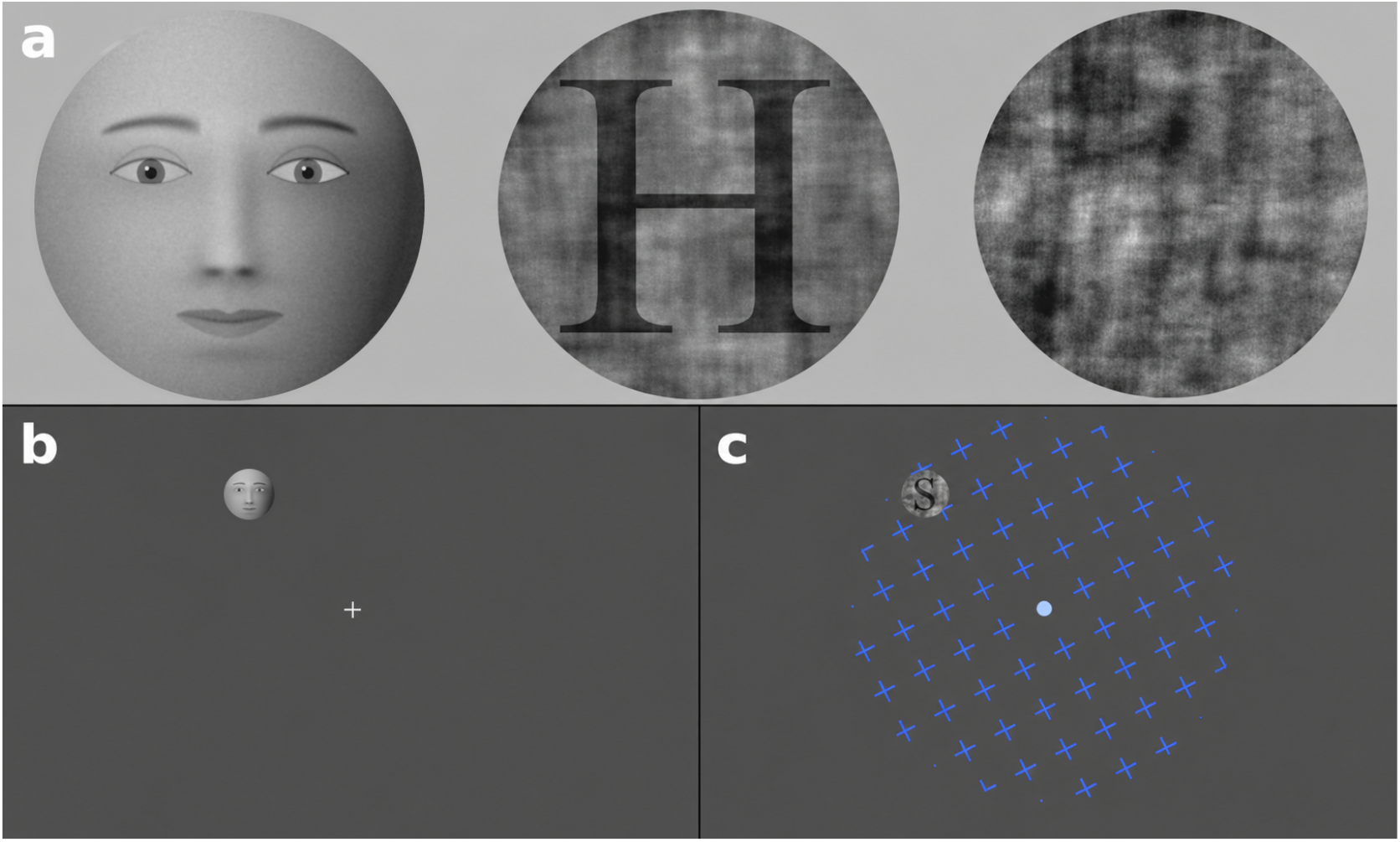
(a) Stimulus examples; (b) Sample screen from Experiment A, used for the peripheral category-discrimination check; (c) Sample screen from Experiment B where a rotating pattern of cross symbols is superimposed on the peripheral target to induce MIB. Note: in panels (a) and (b), the face example is a synthetic, computer-generated face created for illustration only; it does not depict a real person. The actual face stimuli were greyscale photographs of unfamiliar persons from the authors’ internal stimulus database (see Methods, Stimuli); they are not reproduced here in order to protect the identity of the depicted persons.

#### Experiment B

The main experiment applied an MIB paradigm adapted from Sárközy et al. (2024), in which participants fixated a centrally presented blinking fixation spot while a continuously rotating inducer pattern was presented. A rotating (60 deg/s) texture, confined to a circular aperture centred on the fixation spot and spanning 1368 pixels, that is, 16.9 deg in diameter (8.44 deg in radius), composed of + symbols (1 deg size, 0.1 deg stroke RGB [80,100,180]), presented on a uniform grey background (RGB [80,80,80]) was used to induce illusory stimulus disappearances. The direction of rotation was randomly selected as clockwise or anticlockwise. On each trial, a single peripheral target stimulus was presented in the upper-left quadrant, at an effective centre eccentricity of 7.2 deg (583 pixels). The target therefore fell inside the rotating inducer field, close to its outer boundary: the outer edge of the target reached approximately 8.2 deg from fixation (8.3 deg for the largest positional jitter), whereas the inducer field extended to 8.44 deg (684 pixels; Fig. 1c). The central fixation spot had a radius of 0.30 deg (24 pixels). To reduce retinal adaptation, the target position was jittered on each trial in two ways: the polar angle of the target was drawn at random from a uniform range of ±2 deg around the upper-left diagonal, and an additional positional offset of 0.099 deg (8 pixels) was applied in one of five directions (up, down, left, right, or no offset). The resulting centre eccentricity therefore varied between approximately 7.1 and 7.3 deg. The experiment began with a short practice phase, consisting of a single trial, followed by four experimental blocks of 12 trials each (48 trials total), with stimulus category randomized on a trial-by-trial basis within blocks, separated by fixed breaks of 60 s, after which participants self-initiated the next block. Each trial lasted 45 s, during which multiple subjective disappearance and reappearance events occurred. Participants indicated perceived target disappearance by pressing “x” and reappearance by pressing “m”; responses did not alter stimulus presentation and were used solely to record the timing of perceptual reports.

### Stimuli

Stimuli were greyscale images of 16 human faces (8 female) of unfamiliar persons, 16 Roman letters, and 16 Fourier phase-randomized noise images (Fig. 1a; for the face category, a synthetic, computer-generated face is shown in place of the photographs). Face stimuli were drawn from our internally created stimulus database for which consent for scientific use and publication of the stimuli had been obtained. Letter stimuli consisted of uppercase Roman letters [A, B, E, F, G, H, I, J, L, N, P, R, S, T, V, Z], sampled such that the number of letters matched the number of face and noise images. To make the letter stimuli comparable to the other categories in background structure, each letter was superimposed on a phase-randomized background before the final circular cropping and normalization steps. Noise images were generated by randomizing image phase while preserving the Fourier amplitude spectrum of the source face images, thereby retaining their spatial-frequency structure while removing recognizable object content (Näsänen, 1999). All three categories passed through the same final intensity-normalization pipeline. Pixel intensities were standardized for each image according to:

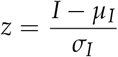

Here, I denotes the original greyscale intensity, μ and σ denote the corresponding image mean and standard deviation, and z denotes the standardized pixel value. The standardized images then underwent the same final rescaling procedure in all three stimulus categories.

In the final stimulus files, pixel values were clipped to the valid 0 to 255 range and the region outside the circular aperture was transparent, so that only the target disc was drawn against the uniform grey background during presentation. Phase randomization and normalization were implemented in MATLAB using custom scripts. Stimulus intensity was quantified as mean greyscale intensity on the final presented target images. Contrast was quantified as mean Weber contrast relative to the uniform background, consistent with the target contrast dimension used in MIB studies (Bonneh et al., 2001; Caetta et al., 2007; Bonneh et al., 2014; Peli, 1990), and as Michelson contrast within the target aperture (Kukkonen et al., 1993). Category values for these and further image measures, computed on the final presented stimuli, are reported in Supplementary Material S1.

To support peripheral discriminability, we first selected a size (diameter: 1.6 deg) that is easily recognizable foveally. Next, we scaled the stimulus size for peripheral presentation using an eccentricity-based linear size increment based on the relationship reported by Anstis (1974). Specifically, target size at eccentricity E (in degrees) was computed as:

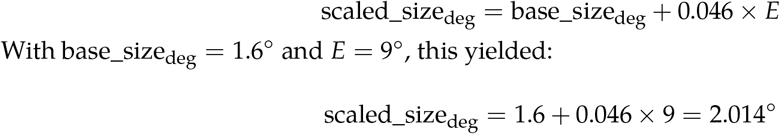

With base_size_deg_ = 1.6° and *E* = 9°, this yielded:

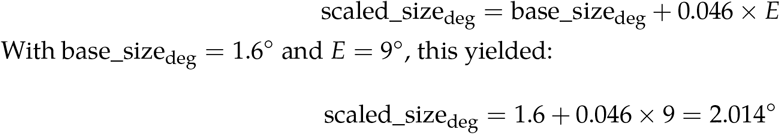

The sizing calculation used the nominal eccentricity parameter of 9 deg in the presentation scripts, which yields 2.014 deg, while the placement routine rendered the target centre at a base eccentricity of 7.2 deg (583 pixels). The final rendered target diameter was 163 pixels, the nearest whole number of pixels, corresponding to 2.012 deg at the conversion factor of 81.00 pixels per degree. Applied at the rendered eccentricity of 7.2 deg, the same size-increment rule would give 1.93 deg, so the target was approximately 4% larger than the size-increment rule prescribes at that eccentricity; this applies identically to all three stimulus categories and therefore does not affect any between-category comparison. Experiments A and B used the same final stimulus set, target size, base upper-left location, viewing distance, and display. Experiment B additionally included the trial-wise positional jitter described above, whereas Experiment A presented the target at the base location. The two experiments also differed in trial duration (3 s versus 45 s) and in the presence of the rotating inducer field. Stimulus statistics reported here and in Supplementary Material S1 were computed from the final rendered stimuli, using the displayed grey background as the intensity reference where appropriate.

### Apparatus

Stimulus generation, including phase randomization and intensity/contrast normalization, was implemented in MATLAB R2023b (The MathWorks, Natick, MA) using custom scripts. Stimulus presentation was implemented in Python 3.9.21 using the Pygame library (version 2.6.1) on a Windows 11 Pro computer. Stimuli were presented on a DELL S2721DGFA monitor (2560 × 1440 pixels; 165.08 Hz; active display width

59.6 cm) at a viewing distance of 105.3 cm. The presentation script converted visual angles to pixels from the full screen width, yielding 81.00 pixels per degree of visual angle; all visual angles reported here are derived from this factor. Head position was stabilized using a chin and forehead rest. A second monitor was used by the experimenter to supervise the experiment. Gaze position was monitored online with an Eye-Link 1000 Plus system (SR Research, Ottawa, Canada) and displayed continuously on the experimenter’s control monitor; participants were re-instructed and re-calibrated when fixation drift was observed. The eye tracker was used for online fixation control only, and gaze samples were not stored for offline analysis. An independent dataset from the same laboratory, recorded on the same tracker at the same target eccentricity with gaze sampled throughout, is reported in Supplementary Material S3.

### Analysis

#### Experiment A

Category discrimination performance was measured as the proportion of correct responses, calculated separately for each stimulus category and across categories. Because noise trials used a no-go response, reaction time was defined only for correct face and letter trials. Participant-level median reaction times for faces and letters were compared using a paired t-test and a Wilcoxon signed-rank test (Student, 1908; Wilcoxon, 1945).

#### Experiment B

Reports of illusory disappearances and reappearances were used to parse trials into epochs of target visibility and invisibility, separately for each trial. For every reported disappearance event, invisible episode durations were defined as the time interval between the disappearance and the subsequent reappearance key presses, yielding a set of invisible episode durations for each trial. If a disappearance was reported without a subsequent reappearance before the end of the trial, the final interval could not be fully observed. In such cases, the duration of this final invisible period was treated as a right-censored observation in the sense of time-to-event analysis (Kleinbaum & Klein, 2012), because the true reappearance time lies beyond the observation window. The observed duration of this interval was computed as the time from the last disappearance to the end of the trial, providing a lower bound on the true invisible episode duration.

This distinction was necessary because trial-level and episode-level analyses have different requirements. For summary measures at the trial level, for example total invisible time and disappearance frequency, censored intervals were included using their observed duration, ensuring that all available information contributed to the overall estimate of perceptual suppression within a trial. In contrast, for duration analyses at the episode level, only completed invisible episodes, in which disappearance was followed by reappearance, were included. The episode-level duration models are therefore conditional on completed episodes: they describe the durations of episodes that both began and ended within a trial rather than of every episode that was initiated, since the true length of a censored episode is unknown.

To quantify the overall strength of the MIB illusion, for each trial we summed the durations of the invisible periods, including censored intervals when present, and normalized this by the total trial duration, yielding an invisibility fraction (IF):

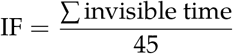

The invisibility fraction ranged from 0 (no disappearance reports) to 1 (target continuously invisible throughout the entire trial). Because multistable phenomena can vary independently in state durations and switching frequency (Leopold & Logothetis, 1999; Brascamp et al., 2005), we quantified disappearance frequency as an onset rate. Specifically, for each trial, we counted the number of disappearance onsets and normalized this count by the total time during which the target was reported as visible, that is, the time window in which a disappearance onset could occur. Visible time was computed as the total trial duration *minus* the total time the target was reported as invisible (including any right-censored interval). The disappearance onset rate was therefore defined as the number of disappearance onsets divided by the total visible time (in seconds), yielding a measure in units of disappearances per second of visible exposure:

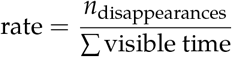

This definition estimates disappearance onsets per unit of time at risk. In the Poisson count model described below, the logarithm of visible time was entered as an exposure offset, following the standard incidence-rate formulation for count data with unequal exposure (Frome, 1983).

Although the sample size for Experiment B was planned a priori for a repeated-measures analysis of variance, that analysis is reported only as a supplementary check: computed on participant-level disappearance rates, it converges with the count model, F(2, 66) = 4.286, p = .0178 (Supplementary Material S2). Experiment B outcomes were analysed at the trial or episode level using models appropriate to the distribution of each measure and the repeated observations within participants. Disappearance counts were analysed with generalized estimating equations (GEE), while transformed continuous outcomes were analysed with participant-random-intercept mixed-effects models (Liang & Zeger, 1986; Laird & Ware, 1982; Pinheiro & Bates, 2000). The specific model for each outcome is described below. Invisibility fraction was modelled on the logit scale:

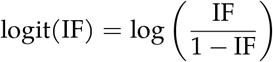

The stimulus category was defined as a categorical fixed effect with noise as the reference level. Block number was entered as a continuous covariate after grand mean centring. For invisibility fraction, values at the boundaries were clipped to [1 × 10^−6^, 1 − 10^−6^] before logit transformation, and the transformed outcome was modelled with a participant random intercept as follows:

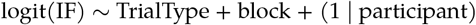

Disappearance frequency was analysed as a count process using a Poisson marginal model (generalized estimating equations; GEE) with clustering by participant. To model the disappearance rate while respecting unequal visible exposure time across trials, we used the log of total visible time per trial as an offset:

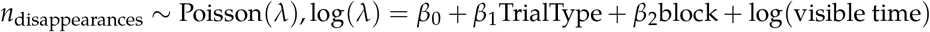

This model yields effect estimates that can be interpreted as incidence rate ratios (exp[β]), reflecting relative differences in disappearance rates between trial conditions and across blocks. An independent working correlation structure was specified, and inference was based on robust sandwich standard errors (Liang & Zeger, 1986). To examine whether stimulus category also influenced the duration of individual invisible episodes, we conducted an event-level mixed-effects analysis. Only completed invisible episodes, in which disappearance was followed by reappearance, were included; right-censored episodes were excluded because their true durations were unknown. For each invisible episode, we also computed the duration of the immediately preceding visible period, from trial onset or the previous reappearance to the current disappearance, as a measure of local temporal context. Invisible episode duration and preceding visible duration were log-transformed to reduce skew and stabilize variance, and preceding visible duration was entered as a covariate because sequential dependencies between successive perceptual episodes are well documented in bistable perception (van Ee, 2009; Pastukhov & Braun, 2011). The model was specified as follows:

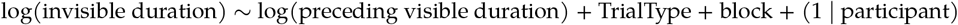

Mixed-effects estimates are reported with Wald-based 95% confidence intervals within the standard mixed-model framework (Laird & Ware, 1982; Pinheiro & Bates, 2000). For the log-link Poisson rate model, coefficients were exponentiated to obtain incidence rate ratios (Frome, 1983). Descriptive summaries were computed at the trial level and after aggregation by participant, and group means are reported with standard errors of the mean (SEM). Within-participant differences were illustrated with paired plots connecting each participant’s category means, and the mean paired difference was reported with a nonparametric bootstrap 95% confidence interval based on 10,000 resamples (Efron, 1979), following an estimation-statistics approach (Ho et al., 2019). Latency to the first disappearance was analysed at the trial level using a Cox proportional hazards model (Cox, 1972), with trials containing no disappearance right-censored at the end of the 45 s trial (Kleinbaum & Klein, 2012). Across 1,632 trials, 1,482 contained at least one disappearance and 150 were censored. Standard errors for this model were clustered by participant, so that the repeated trials contributed by the same observer were treated as one cluster (Lin & Wei, 1989). Visible episode durations preceding disappearance onsets were analysed as an exploratory outcome in an episode-level mixed-effects model with log visible duration as the outcome, stimulus category with noise as the reference and centred block number as fixed effects, and a participant random intercept. Unlike the invisible episode model, this model included no covariate for the duration of the preceding episode. Trial type was coded with noise as the reference category, yielding the two planned contrasts of letter versus noise and face versus noise; direct face-versus-letter contrasts are reported as exploratory. No multiplicity correction was applied to these contrasts, which are two model-based comparisons against a common reference rather than a full pairwise family. Supplementary Material S2 reports Holm-adjusted values for these contrasts (Holm, 1979), together with descriptive distributions and robustness analyses.

Analysis was performed in Python 3.13.9, using the *statsmodels, NumPy, pandas*, and *Matplotlib* libraries.

## Results

### Experiment A

Overall category-discrimination accuracy for the peripheral stimulus was 85.93% (580/675 correct; Wilson score 95% CI [83.10%, 88.35%]; Wilson, 1927), with participant-level accuracy ranging from 72.59% to 92.59%. Accuracy by category was 93.33% for faces, 88.00% for letters, and 76.44% for noise images. Median reaction times on correct go trials were 693 ms for faces and 745 ms for letters; the participant-level medians did not differ reliably (paired t(4) = −1.56, p = .194; Wilcoxon p = .312).

### Experiment B

#### Higher disappearance rates for faces and letters than for noise

To test category effects on switching frequency while controlling for unequal visible exposure times, disappearance counts were analysed using a Poisson GEE with log(visible time) as an offset. Relative to noise, disappearance rates were higher for faces (IRR = 1.090, 95% CI [1.027, 1.157], p = .005) and for letters (IRR = 1.044, 95% CI [1.004, 1.087], p = .031), as shown in Fig. 2a. Consistent with these model estimates, participant-level mean disappearance rates were lowest for noise (M = 0.149, SEM = 0.016), intermediate for letters (M = 0.154, SEM = 0.016), and highest for faces (M = 0.161, SEM = 0.016; Fig. 2b). The exploratory direct face-versus-letter contrast was not reliable (IRR = 1.043, 95% CI [0.987, 1.103], p = .132).

**Fig. 2.**
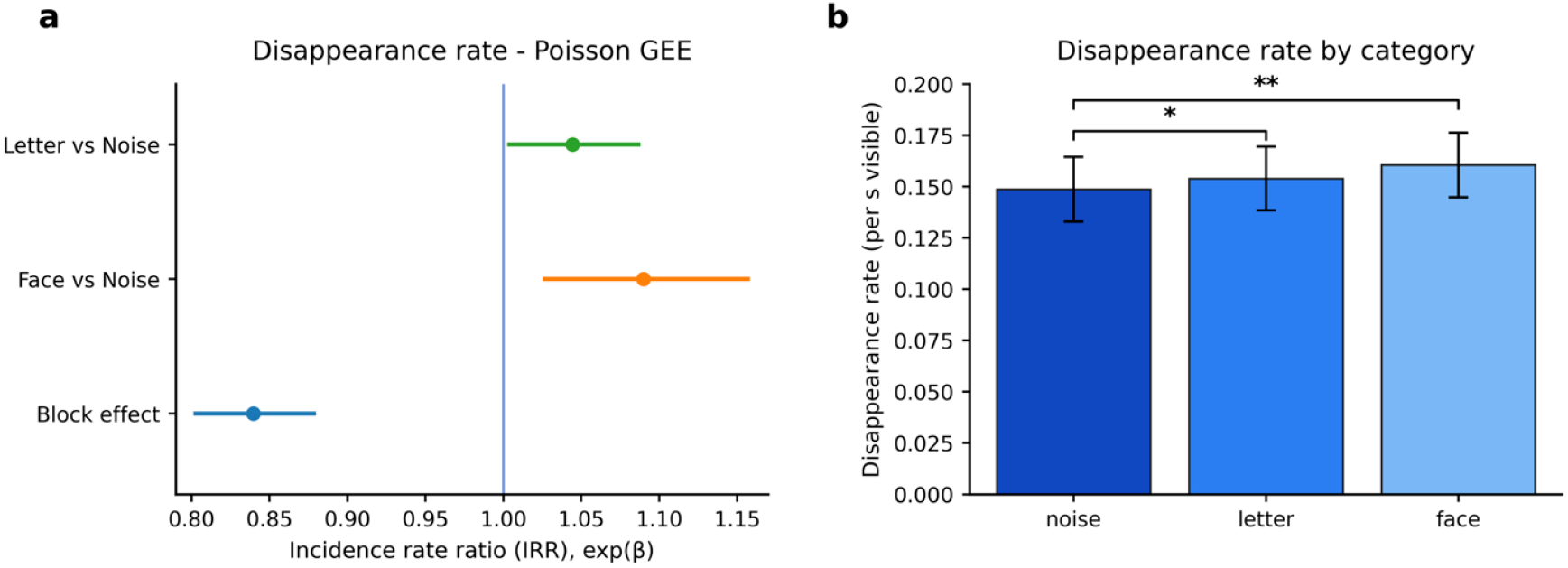
Disappearance rate by stimulus category and block. (a) Forest plot of Poisson GEE fixed effects, expressed as incidence rate ratios (IRRs) with 95% confidence intervals. Values above 1 indicate higher disappearance rates relative to the noise reference condition, whereas values below 1 indicate lower rates. The block effect indicates the multiplicative change in disappearance rate across successive blocks. (b) Participant-level mean disappearance rates for each stimulus category, with error bars indicating SEM. Asterisks denote the significance of the corresponding model-based contrast against the noise reference condition (* p < .05; ** p < .01; *** p < .001).

To illustrate the size of these category effects within participants, we also plotted paired differences across participants in disappearance rates. On average, participants showed a higher disappearance rate for faces than for noise stimuli, with a mean paired difference of 0.012 disappearances/s visible (95% bootstrap CI [0.003, 0.021]; Fig. 3a). For letters, relative to noise, the mean paired difference was 0.005 disappearances/s visible (95% bootstrap CI [−0.0006, 0.011]; Fig. 3b) suggesting a smaller and less consistent participant-level difference than for faces. These paired plots are included to show the size and direction of the within-participant differences. The primary inferential analysis was the Poisson GEE, which showed higher disappearance rates for both faces and letters than for noise (p = .005 and p = .031, respectively).

**Fig. 3.**
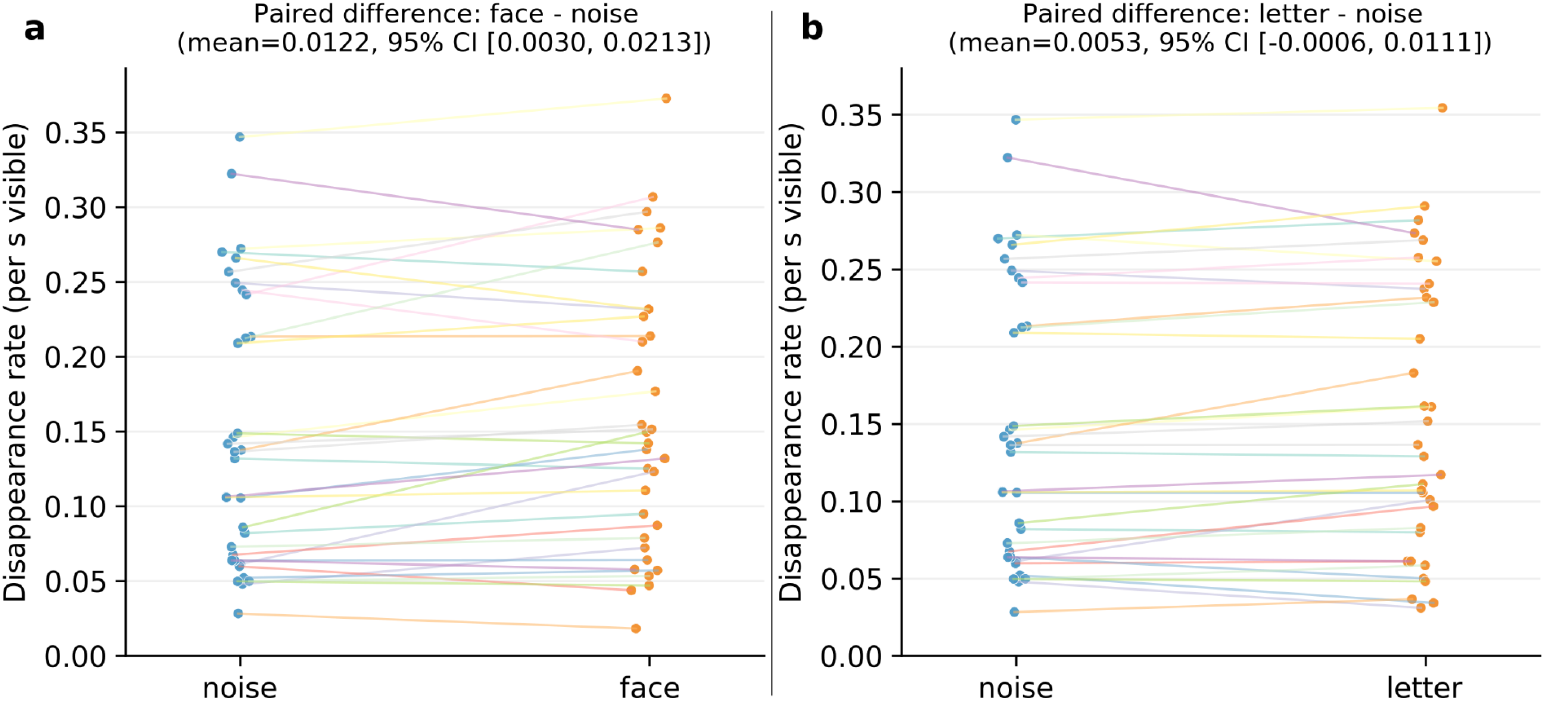
Paired plots of participant-level disappearance rates. (a) Face versus noise disappearance rates, with each line connecting paired participant means. The title reports the mean paired difference, face minus noise, with its bootstrap 95% confidence interval. (b) Letter versus noise disappearance rates, plotted in the same format. The title reports the mean paired difference, letter minus noise, with its bootstrap 95% confidence interval.

#### Shorter invisible episodes for face stimuli

To determine whether category effects were also expressed in the duration of individual invisible episodes, an episode-level mixed-effects model of log invisible duration included log preceding visible duration, trial condition with noise as the reference category, centred block number, and a participant random intercept. Analyses were performed on 9,090 completed invisible episodes, with censored tails excluded. Invisible episodes were shorter for faces than for noise images (β = −0.046, 95% CI [−0.078, −0.015], p = .004), corresponding to a duration multiplier of exp(β) = 0.955. In contrast, invisible episode durations for letters did not differ from noise (β = −0.001, 95% CI [−0.032, 0.031], p = .974; Fig. 4a; model-adjusted geometric means are shown in Fig. 4b). The exploratory direct face-versus-letter contrast was β = −0.046, 95% CI [−0.077, −0.015], p = .004, corresponding to a duration multiplier of 0.955; it is numerically almost identical to the face-versus-noise contrast because letters did not differ from noise. A participant-clustered Gaussian GEE sensitivity analysis likewise retained the face-versus-noise effect (β = −0.047, 95% CI [−0.084, −0.009], p = .016; Supplementary Material S2).

**Fig. 4.**
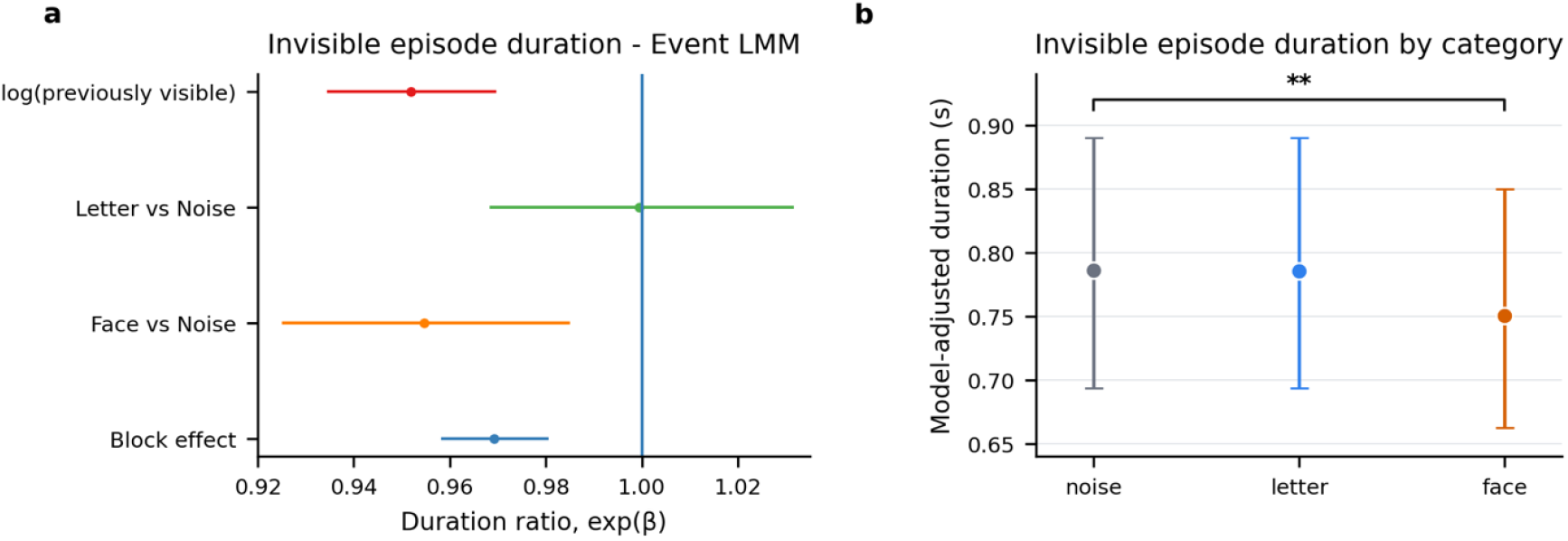
Invisible episode duration. (a) Forest plot of the event-level model fixed effects on invisible episode duration. Because the model was fitted to log-transformed durations, coefficients are shown after exponentiation as duration ratios with 95% confidence intervals. Values below 1 indicate shorter invisible episodes and values above 1 indicate longer invisible episodes. Trial type effects are expressed relative to the noise reference condition. The block effect represents the multiplicative change per successive block, and log(previously visible) represents the association between the duration of the preceding visible period and the subsequent invisible episode. (b) Model-adjusted geometric mean invisible episode duration for each stimulus category, back-transformed from the log scale, with 95% confidence intervals, plotted on an axis scaled to the range of the estimates so that the direction and the size of the category difference are visible. Asterisks denote the significance of the corresponding model-based contrast against the noise reference condition (* p < .05; ** p < .01; *** p < .001). Raw distributions of completed invisible and visible episode durations are shown in Supplementary Material S2, Figure S2.1.

#### Visible episode durations

Visible episode durations preceding disappearance onsets were analysed as an exploratory outcome with the episode-level model specified in the Analysis section, on 9,330 visible episodes. Relative to noise, visible episodes were shorter for faces (β = −0.053, 95% CI [−0.087, −0.018], p = .003; duration multiplier = 0.949), whereas the corresponding effect for letters was smaller and did not reach significance (β = −0.033, 95% CI [−0.068, 0.001], p = .059). Visible episodes also became longer across successive blocks (β = 0.102, 95% CI [0.089, 0.114], p < .001), mirroring the decrease in disappearance rate. Mean visible episode durations were 5.33 s for noise, 5.17 s for letters, and 5.09 s for faces, with medians of 3.61, 3.47, and 3.49 s, respectively. Because the disappearance rate was expressed per second of visible time, it is closely related by construction to the mean duration of visible episodes, so these two outcomes are not independent confirmations of one another; the outcomes that constrain the interpretation independently are invisible episode duration and overall invisibility fraction. Together with the invisible episode results, the shorter visible episodes for faces indicate a faster cycle of perceptual alternation rather than a selective shortening of the suppressed state.

#### Block number and the duration of preceding visible periods

We also examined whether disappearance dynamics changed over the course of the experiment and whether they were related to the duration of the immediately preceding visible period. To capture potential changes over time, we included block number as a predictor, where each block represents a successive segment of the experiment (blocks 1–4). Disappearance rate was systematically associated with block number, decreasing across successive blocks (IRR = 0.84 per unit increase in centred block, 95% CI [0.802, 0.878], p < .001; Fig. 2a). This indicates that disappearances occurred less frequently as the experiment progressed. A similar pattern was observed for overall invisibility fraction, which was highest in the first block and lower in subsequent blocks (Fig. 5). Because block number indexes time on task rather than a manipulated factor, we treat this association as descriptive.

**Fig. 5.**
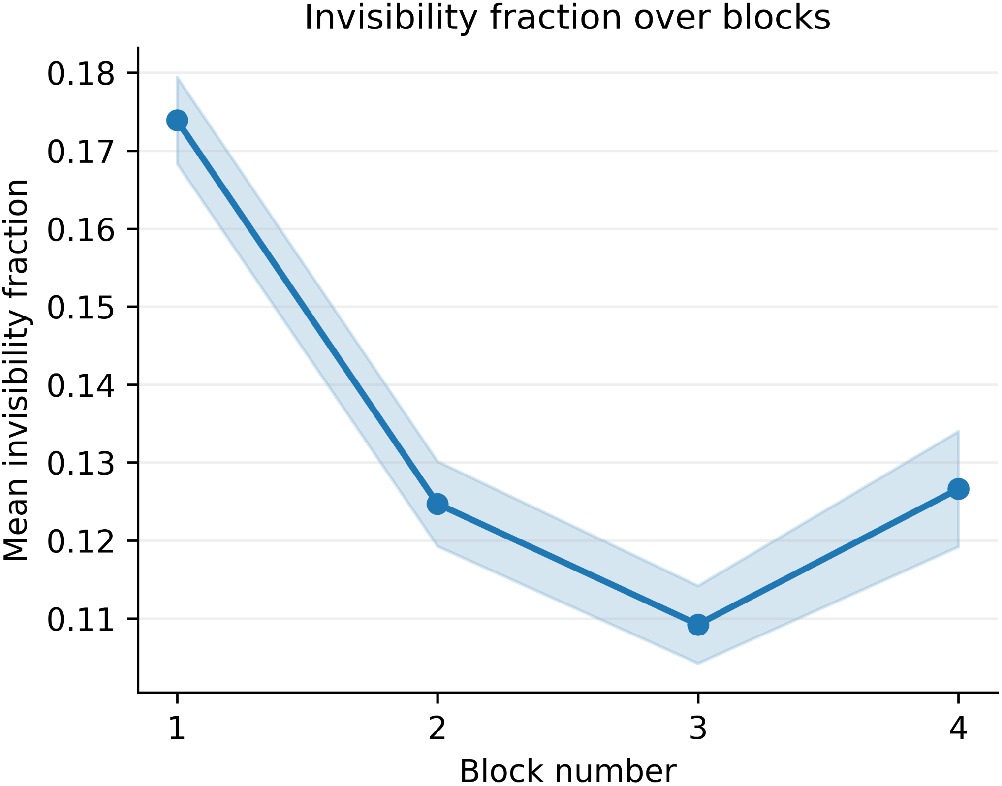
Mean invisibility fraction across blocks. The shaded band gives ±1 standard error of the mean computed across all trials within a block. Because trials are nested within participants, this band describes the precision of the trial-level mean rather than the spread of the individual trials, and it is not a participant-level standard error; the inferential estimate of the block effect comes from the participant random intercept model reported in the text.

Independent of target stimulus category, the duration of a visible period predicted the duration of the subsequent invisible period (β = −0.049, 95% CI [−0.067, −0.032], p < .001), indicating that longer visible intervals tended to be followed by shorter invisible episodes. Invisible episodes were also shorter in later blocks (β = −0.031, 95% CI [−0.044, −0.019], p < .001), consistent with the event-level duration ratio estimate shown in Fig. 4a.

#### Overall invisibility fraction did not differ reliably by category

At the trial level, invisibility fraction was analysed with a logit-transformed mixed-effects model with a participant random intercept, with stimulus category, using noise as the reference, and centred block number entered as fixed effects. There was a robust block-related reduction in invisibility fraction (β = −0.567, 95% CI [−0.689, −0.446], p < .001), consistent with the descriptive means across blocks 1 to 4 (0.174, 0.125, 0.109, 0.126; Fig. 5). In contrast, invisibility fraction did not differ reliably by stimulus category: letter versus noise β = 0.225, 95% CI [−0.107, 0.558], p = .184; face versus noise β = 0.229, 95% CI [−0.104, 0.561], p = .178. The descriptive means were 0.132, 0.134, and 0.135 for noise, letters, and faces, respectively. A boundary-compatible fractional-response GEE sensitivity analysis that included zero values without clipping likewise showed no reliable category effect (letters versus noise β = 0.020, 95% CI [−0.038, 0.079], p = .495; faces versus noise β = 0.030, 95% CI [−0.053, 0.113], p = .482; Supplementary Material S2).

#### Latency to first disappearance did not differ reliably by category

In the Cox model, using noise as the reference condition, stimulus category did not reliably affect the timing of the first disappearance (Fig. 6a–b). Results are expressed as hazard ratios, which reflect the relative likelihood that a disappearance occurs at a given moment compared to the reference condition. There was no reliable difference between letters and noise (HR = 1.064, 95% CI [0.961, 1.178], p = .235), nor between faces and noise (HR = 0.994, 95% CI [0.887, 1.113], p = .918). In contrast, block number had a strong effect: the hazard ratio decreased across blocks (HR = 0.783 per unit increase in centred block, 95% CI [0.732, 0.837], p < .001), indicating that first disappearances tended to occur later as the experiment progressed (Fig. 6b).

**Fig. 6.**
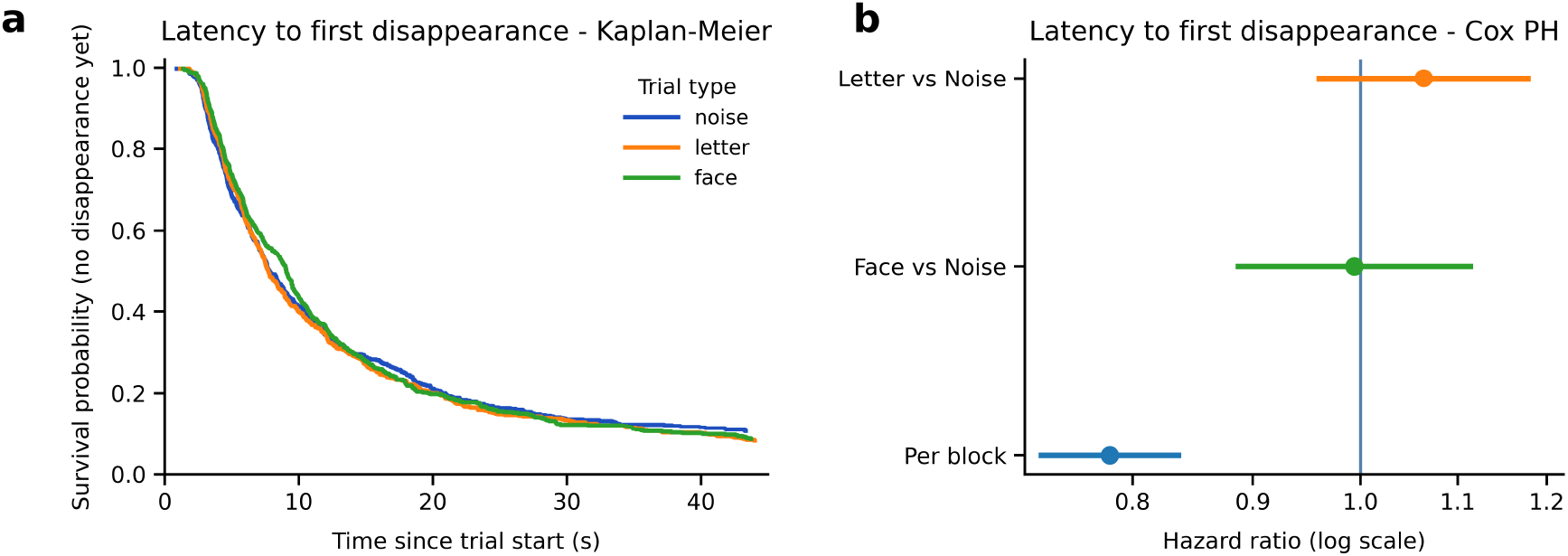
Latency to first disappearance. (a) On the Kaplan–Meier survival plot, the y-axis gives the probability that no first disappearance has occurred yet over time; lower/steeper curves indicate earlier onsets (higher hazard). The close overlap of curves indicates highly similar latency distributions across stimulus categories. (b) On the Cox hazard-ratio forest plot, points show hazard ratios (HRs) with 95% CIs around the HR = 1 “no-effect” line; HR > 1 indicates earlier onsets and HR < 1 later onsets. CIs crossing 1 indicate no reliable effect.

## Discussion

The current study tested whether stimulus category is associated with the temporal dynamics of motion-induced blindness. Using human faces, familiar Roman letters, and phase-randomized noise images as peripheral targets, we quantified disappearance rate, invisible episode duration, visible episode duration, overall invisibility fraction, and latency to first disappearance. The results reveal category-related differences in the temporal organization of MIB, with the clearest and most consistent effects observed for faces relative to the noise baseline.

### Category-related differences in the dynamics of perceptual alternation

The clearest category effect was observed in disappearance rate. Relative to the phase-randomized noise baseline, disappearance rates were higher for faces and, to a smaller extent, for letters. Faces also showed shorter invisible episodes than noise. Together, these findings indicate that stimulus category is associated with how frequently perceptual transitions occur and how long suppression episodes persist. Considered alone, the combination of more frequent and shorter invisible episodes for faces could suggest prioritized access to awareness (Lanfranco et al., 2023). The complementary analysis of visible episodes, however, shows that the temporal effect is broader than a selective shortening of suppression.

These findings are in line with previous work suggesting that category-related information remains functionally relevant even during periods without conscious access (Harris et al., 2013) and that object representations can be updated in the absence of awareness (Mitroff & Scholl, 2005). The present results extend this literature by showing that target category is associated with the temporal organization of alternations between visible and invisible states, even though the overall invisibility fraction did not differ reliably. Experiment A further indicated that the stimulus categories were discriminable at the target size and base peripheral location used in Experiment B, under static, no-inducer conditions.

The pattern across dependent measures helps characterize the nature of this modulation. Faces showed a higher disappearance rate and shorter invisible episodes, while the overall invisibility fraction and the latency of the first disappearance did not differ reliably across categories. This indicates that the category effect was not expressed simply as a uniform increase in the amount of suppression. Instead, it was expressed in the timing of perceptual transitions. Such a pattern is compatible with the possibility that category-level information about the target remains available during suppression and contributes to the temporal organization of perceptual alternations (Mitroff & Scholl, 2005; Harris et al., 2013), although the present design does not identify the specific mechanism responsible for that modulation.

The analysis of visible episodes provides complementary information about whether the category effect is specific to the suppressed state. A recent study comparing breaking and reverse-breaking CFS found that the face-inversion effect was larger for breaking suppression than for reverse-breaking suppression (Ciorli et al., 2025). In the present data, faces showed shorter invisible and visible episodes of comparable relative magnitude, while the overall invisibility fraction showed no reliable category difference. Thus, the category-related effect is better described as a change in the tempo of perceptual alternation than as a selective shortening of suppression. Within a dynamical account of bistable perception, this pattern is consistent with altered transition dynamics in both directions, with no reliable difference in the overall balance between perceptual states.

### No reliable category effects on overall invisibility fraction and onset latency

Despite significant category differences in disappearance rate and invisible episode duration, overall invisibility fractions and the hazard of the first disappearance showed no reliable effect of stimulus category. This dissociation highlights the multidimensional nature of bistable perceptual processes, whereby disappearance rate, state duration, and cumulative dominance time capture partly dissociable dimensions of perceptual dynamics (Leopold & Logothetis, 1999; Brascamp et al., 2005). The observed near-equality of the invisibility fractions is consistent with the symmetry of the duration effects: for faces a higher rate of disappearance was accompanied by shorter episodes of both kinds, so that the proportion of time suppressed showed no reliable change. The absence of a reliable category difference in the latency of the first disappearance does not support a general readiness to report faces as disappearing earlier. It also indicates that the initial transition into suppression and the subsequent alternation between the two states need not be modulated in the same way, although the present design does not identify what distinguishes them.

### Integration with prior literature on attentional modulation and category-specificity

The result that faces disappeared more often might seem counterintuitive, given their known attentional priority (Bindemann et al., 2005; Palermo & Rhodes, 2007). However, such seemingly counterintuitive results have been found in MIB before: the allocation of spatial attention to the target can increase the probability of disappearance (Schölvinck & Rees, 2009), possibly because of the enhanced neural competition at the object-mask boundary. One possible interpretation is that faces engage attention more strongly than other categories, and this paradoxically increases the probability of perceptual suppression, consistent with models of response gain, where attention increases both signal and competition (Carrasco, 2011; Ling & Carrasco, 2006).

One possible interpretation of the present pattern is that the high attentional salience of faces is associated with a higher number of suppression events and a shorter persistence of each event, perhaps because of rapid re-engagement of category-specific processing streams. The present design contains no independent measure of attentional engagement or category-selective cortical activity, so this remains a hypothesis. Letters, being familiar and highly overlearned, may engage attention and category-specific processing in related ways, but the effects observed for letters were smaller and less consistent. These findings are compatible with electrophysiological evidence that category-related signals can be maintained during periods of conscious suppression without requiring a fresh onset response upon reappearance (Harris et al., 2013), suggesting that perceptual transitions may be better characterized by changes in access to ongoing representations than by their reinitialization.

### Implications for theories of perceptual awareness

The present pattern extends object-level accounts of MIB, according to which suppression reflects altered conscious access to an ongoing object representation rather than the complete absence of that representation from visual processing (Mitroff & Scholl, 2005; Harris et al., 2013). With size and position matched and measured intensity and contrast closely comparable, the association between stimulus category and temporal dynamics is consistent with the view that higher-level representations can contribute to the organization of conscious access. This interpretation is also compatible with models that describe conscious perception as emerging from interactions between feedforward and recurrent processing across representational hierarchies (Lamme, 2006; Dehaene & Changeux, 2011). Related low-versus high-level questions have shaped binocular rivalry and continuous flash suppression research for many years (Tong, Meng, & Blake, 2006; Hesse & Tsao, 2020; Ciorli et al., 2025). The present results suggest that analogous category-related effects can also be observed during motion-induced suppression without interocular competition, although whether this generalizes across multistable paradigms remains to be established.

Our results are also compatible with recent dynamical accounts of MIB and bistable perception in general. MIB has been described as a noisy excitable system in which perceptual transitions emerge from the interaction of adaptation, competition, and stochastic fluctuations (Katkov et al., 2024). Within such a framework, differences related to stimulus category could arise if stimulus properties at higher levels of representation modulate system parameters such as excitability, baseline input, or transition threshold, thereby biasing switching behaviour. More generally, perceptual bistability has been characterized as the behaviour of neural systems operating near criticality, where small fluctuations can trigger transitions between perceptual states (Freyer et al., 2011; Cocchi et al., 2017). Empirical work identifying critical slowing down and noise-induced intermittency in bistable perception further supports this view (Pisarchik et al., 2014). Although the present data do not directly test these mechanisms, they are consistent with the idea that category-related signals can shape the dynamical conditions under which conscious access fluctuates.

Two further, category-independent features of the present data deserve comment. First, the disappearance rate decreased systematically across successive blocks. Several non-mutually-exclusive mechanisms could contribute to this. Neural adaptation is a well-supported contributor to MIB (Gorea & Caetta, 2009), and prolonged exposure could, in addition, reduce the efficacy of the moving mask across blocks. A reduction in attentional engagement with time on task is also plausible, since withdrawing attention from the target reduces the frequency of perceptual transitions (Schölvinck & Rees, 2009; Geng et al., 2007); however, this account predicts longer invisible episodes, whereas we observed shorter ones in later blocks. Perceptual learning of the target-mask configuration, used here in the general sense of experience-dependent improvement or stabilization in visual processing (Sagi, 2011), is compatible with the observed joint decrease in disappearance rate and invisible episode duration and with the lengthening of visible episodes. Because block number indexes time on task, these candidate mechanisms cannot be distinguished in the present design. Second, the negative relationship between the duration of a visible episode and that of the subsequent invisible episode is consistent with a system in which perceptual alternations are shaped by the interplay of adaptation and noise: a long dominance period allows adaptation of the dominant representation to accumulate, which can influence the ensuing suppressed period. Serial dependencies of this kind are documented in bistable perception (van Ee, 2009; Pastukhov & Braun, 2011) and are compatible with dynamical accounts in which MIB is treated as a noisy excitable system (Katkov et al., 2024).

### Limitations and future directions

Several limitations should be considered. The stimulus categories differed in structural properties beyond their categorical identity, including shape complexity and local feature structure, which the phase-randomized controls constrain for the face-versus-noise comparison without eliminating. The study restricted targets to the upper-left quadrant, which limits spatial generalizability; given known anisotropies in MIB across the visual field (Sárközy et al., 2024), future studies should test whether category-related effects generalize across target locations, including locations relevant to hemispheric asymmetries in face and letter processing (Rossion et al., 2003; Cohen et al., 2000). Perceptual transitions were measured by manual report, which introduces response latency and variability; objective or neural measures could complement self-report in future work (Donner et al., 2008; Schölvinck & Rees, 2010). Natural images are in general more appropriately compared with other natural images than with artificial stimuli, so objects or scenes would be the more suitable comparison category for a follow-up experiment. Additional target categories, including natural non-face categories such as objects or scenes, and manipulations of attentional load would help clarify the relative contributions of semantic salience, cortical specialization, visual expertise, and attention to MIB dynamics (Schölvinck & Rees, 2009). Identifying which specific image property generates a category difference would require parametric manipulation of features within a category, which the present design does not attempt.

Physical equivalence of complex stimulus sets cannot be established exhaustively by any single scalar descriptor. The intensity and contrast measures reported here were closely comparable across categories, while residual differences in higher-order image structure remained, including a small orientation bias: relative to the noise baseline, the letter set carried an approximately 7% stronger cardinal-to-oblique amplitude ratio, in the direction that the predominantly vertical and horizontal strokes of Roman characters imply. This difference is small relative to the variation between individual exemplars within each set, and it is reported descriptively; no analysis in the present study depends on it. To place the measured residuals on a scale, the published contrast dependence of MIB (Bonneh et al., 2014) implies that contrast differences of approximately 33% for faces and 16% for letters would be required to produce the observed disappearance-rate ratios, whereas the measured differences in mean Weber contrast were +1.80% and −0.84%; on that relation neither residual predicts a rate change larger than about 2% in magnitude, and the prediction for letters carries the opposite sign to the observed effect. The full quantitative verification of the presented stimuli, comprising the intensity and contrast measures, the exemplar-level correspondence between the letter and noise sets, the rotationally averaged amplitude spectra and the orientation summaries, is reported in Supplementary Material S1, from which these properties can be checked directly. Future work using stimulus sets prospectively equated within the visible aperture would provide a still stronger physical control.

The eye tracker was used for online fixation control only and gaze samples were not stored, so fixation stability and microsaccade dynamics could not be quantified offline. Microsaccade rate decreases before an illusory disappearance and increases before a reappearance, so that fixational eye movements bear opposite-signed relationships to the two perceptual transitions (Bonneh et al., 2010; Hsieh & Tse, 2009). A category difference confined to the overall rate of fixational eye movements would therefore be expected to move disappearance frequency and invisible episode duration in the same direction, predicting either fewer and shorter episodes or more frequent and longer ones. Faces showed more frequent but shorter episodes, which is neither pattern. Because no gaze record of Experiment B exists, we additionally analysed an independent dataset from the same laboratory in which gaze was recorded throughout, reported in Supplementary Material S3. It comes from a motion-induced blindness experiment run on the same eye tracker, at the same target eccentricity and within an inducer field of nearly the same size, and three of its five target classes were phase-randomized control images, faces and Roman letters; across 2,674 trials from 35 participants, fixation stability, gaze position, EyeLink EBLINK event-record rate, EyeLink ESACC event-record rate and saccade amplitude showed no reliable differences between those three classes. The only omnibus test below .05 before correction concerned the proportion of usable gaze samples rather than eye movements, and it did not survive correction. In that dataset the behavioural category effect was itself absent, and its design differed from Experiment B in several ways that could have reduced sensitivity to the category question, with shorter trials, four target locations, a smaller target whose peripheral discriminability was not separately assessed, and behavioural values that remained descriptively lower after recomputation over the response-enabled window; Supplementary Material S3, Section 2 sets out that comparison, and the dataset is therefore reported as a control on the stimulus classes under a closely related but not identical MIB configuration rather than as evidence for or against the effects reported here. Supplementary Material S3 reports this control in detail, including the trial-by-trial linkage and quality control, three fixation-window definitions, participant-level effect sizes with bootstrap confidence intervals for every oculomotor measure, the exploratory pupil analyses, and the four limits of the conclusion that the dataset supports. These are not the eye movements of Experiment B, and the inference is correspondingly indirect, but they provide no evidence for the version of the confound in which these stimulus classes elicit different tonic oculomotor behaviour under the conditions of that dataset. This argument constrains only a tonic difference in microsaccade rate; it does not extend to transition-locked, directional or amplitude differences, and an oculomotor contribution to the present category effects therefore cannot be excluded. Future studies should record gaze continuously; pupillometry would provide a complementary index of stimulus processing.

Because MIB provides no physically defined event corresponding to an illusory disappearance, standard signal detection analysis cannot be applied directly to the disappearance reports themselves. Physical replay, no-report measures, or additional psychophysical probes of sensitivity and criterion would allow perceptual changes to be separated more directly from category-dependent report criteria (Caetta et al., 2007; Bonneh et al., 2010; Tsuchiya et al., 2015; Hesse & Tsao, 2020). None of these was included in the current study, so a category-dependent shift in the readiness to report perceptual transitions cannot be excluded as a contribution to the observed effects.

Sample composition limits generalizability. Experiment B included 29 female and 5 male participants, and Experiment A included female participants only. Generalization to a more balanced sample remains to be established.

## Conclusion

The present study shows that target category is associated with the temporal dynamics of motion-induced blindness. Faces produced more frequent disappearances together with shorter visible and invisible episodes, whereas the overall fraction of time reported invisible and the latency to first disappearance were not reliably altered. This pattern is best described as a change in the tempo of perceptual alternation rather than a selective change in suppression depth or entry into awareness. Together, the findings extend category-related effects in perceptual suppression to MIB and provide a basis for testing how higher-level stimulus representations interact with the dynamics of conscious access. Future work combining tighter stimulus matching with replay or no-report measures can determine the relative contributions of perceptual and decisional mechanisms.

## Acknowledgements

We thank Jakob Hohwy for his comments on the paradigm. We also gratefully acknowledge the support of the INTREPID Consortium and the broader Accelerating Research on Consciousness initiative, which promotes open science and adversarial collaboration in consciousness research (https://www.templetonworldcharity.org/blog/integrated-i nformation-theory-iit-and-predictive-processing-accounts-of-consciousness).

## AI-use disclosure

AI-based tools were used for proofreading, consistency checking across documents, and independent cross-checking of the stimulus and analysis scripts. All numerical results and stimulus measurements reported here were independently recomputed from the underlying data and stimulus files, and all scientific interpretations and editorial decisions were made and approved by the authors, who take full responsibility for the final manuscript.

## Data Availability

The raw behavioural data of Experiments A and B supporting the findings of this study are available on the Open Science Framework at https://osf.io/xvefm. The repository additionally contains the original stimulus set, the presentation scripts of Experiments A and B, the results, and the analysis and figure scripts. Supplementary Material S3 reports the independent eye-tracking control dataset; its stimulus set, processed data, results, and analysis and figure code are deposited in the same repository, in the folder MIB_eye_tracking_control_experiment_analysis, and its raw EyeLink EDF recordings are available from the authors on reasonable request. Supplementary Materials S1, S2 and S3 accompany the manuscript.

## Author Contributions

András Sárközy: Conceptualization, Methodology, Software, Formal analysis, Investigation, Data curation, Visualization, Writing – original draft, Writing – review and editing.

Sophia Gommlich: Methodology, Investigation, Resources, Data curation, Writing – original draft, Writing – review and editing.

Jonathan E. Robinson: Conceptualization, Methodology, Writing – original draft, Writing – review and editing.

Gyula Kovács: Conceptualization, Methodology, Supervision, Project administration, Funding acquisition, Formal analysis, Writing – original draft, Writing – review and editing.

All authors critically reviewed and approved the final manuscript for submission.

