## Supplementary_Material_1 for "Stimulus category is associated with conscious access during motion-induced blindness"

Quantitative verification of the low-level image properties of the stimulus set

### 1. Stimuli

The stimulus files are 420 × 420 pixel 8-bit greyscale PNGs with a binary alpha channel that defines a hard-edged circular aperture of 210 pixel radius. The region outside the aperture is fully transparent and was never visible on screen. The presentation script resampled each image to 163 × 163 pixels with pygame.transform.scale, the unfiltered fast scale of the Pygame library, which at the conversion factor of 81.00 pixels per degree used throughout corresponds to a rendered target diameter of 2.012 deg.

With the single exception noted below, all analyses reported here were carried out on the image as the observer received it, that is, resampled to 163 × 163 pixels by the same rule the presentation routine applied and composited onto the uniform grey field (RGB [80,80,80]) against which the target was displayed. That rule is unfiltered nearest-neighbour sampling with source index floor(i × 420 / 163), which is what pygame.transform.scale computes, and the analysis script reproduces it exactly. Pixels outside the circular aperture were excluded from every intensity and contrast measure. The exception is the exemplar correspondence of Section 3, which is recovered from the native 420 × 420 files, because the finer sampling separates the candidate pairings more sharply. As reported in Section 3, the presented images recover the same correspondence.

Each measure was computed separately for each of the 48 images. Because the three sets are a fixed stimulus inventory rather than a sample drawn from a population of images, the primary presentation is descriptive: category means with the spread across exemplars are given in Table S1.1 and the per-image values are shown in Figure S1.1. The one comparison for which a paired analysis is defined is the letter versus noise comparison, for the reason given in Section 3.

**Reproducibility.** Every number, table and figure in this supplement is produced by a single script, S1_lowlevel_analysis.py, which is archived with the 48 stimulus files in the study repository. The script documents the resampling rule, the contrast map definition, the treatment of the region outside the aperture, the FFT normalization, the radial binning, the orientation binning and the conversion to cycles per degree, so that the verification reported here can be regenerated independently. Fourier amplitudes are unnormalized magnitudes; every spectral quantity reported below is either a ratio between categories or a share within a single image, so any constant normalization factor cancels. The only quantities in this supplement that the script does not derive from the stimulus files themselves are the two observed disappearance rate ratios used as inputs to the benchmark of Section 4; these come from the primary behavioural model and are supplied to the script as clearly labelled constants. One choice materially affects the spectral results. The Fourier analysis is applied to the contrast image $c(x,y) = (V(x,y) - B)/B$ inside the aperture and zero outside, that is, to the display-input contrast map presented within the aperture, and not to the raw file. Because the display was not photometrically calibrated, this is a contrast map of 8-bit code values and not of measured luminance. Taking the FFT of the raw file instead would include the invisible surround, whose hard circular edge dominates the spectrum identically in all three categories and therefore artificially compresses the differences between them.

Because the display was not photometrically calibrated, all values reported here are 8-bit greyscale drive values and not measured cd/m². We therefore use the term greyscale intensity rather than luminance throughout.

### 2. Intensity and contrast measures

Two definitions of contrast are reported, one referencing the background and one referencing the target itself. With V the 8-bit pixel value inside the aperture, μ its mean, Vmax and Vmin its extreme values, and B = 80 the background value: mean Weber contrast is $(\mu - B)/B$, and Michelson contrast within the target is $(V_{max} - V_{min})/(V_{max} + V_{min})$.

Mean Weber contrast is our primary measure, because it is the dimension along which the contrast dependence of MIB has actually been parameterized, for targets that were uniform patches or Gaussian luminance blobs (Bonneh et al., 2001; Hsu et al., 2004; Caetta et al., 2007; Gorea & Caetta, 2009; Bonneh et al., 2014), and because for a patch presented on a uniform field of a different intensity it is the background rather than the mean of the patch that constitutes the adapting level. Michelson contrast within the target is reported alongside it because it summarizes the intensity range the target itself carries (Kukkonen et al., 1993), and because no single scalar definition of contrast is adequate for complex images (Peli, 1990); the spatial structure that a scalar measure cannot capture is treated separately in Section 5. Because Michelson contrast is defined on the extreme pixel values inside the aperture, it is the more sensitive of the two measures to the resampling rule. The script therefore recomputes both measures under four alternative interpolation rules as a check. Mean Weber contrast is essentially unaffected by that choice, the difference from the noise baseline remaining between −0.84% and −1.04% for letters and between +1.59% and +1.80% for faces; the Michelson difference varies more, and Section 4 shows that the conclusion drawn there holds under every rule tested.

**Table S1.1.** Mean greyscale intensity, mean Weber contrast relative to the background, and Michelson contrast within the target, by stimulus category (n = 16 images per category; SD across images in parentheses). The last two columns give the difference from the noise baseline.

| Measure | Noise | Letters | Faces | Letters vs noise | Faces vs noise |
|---|---|---|---|---|---|
| Mean greyscale intensity (8-bit) | 94.3276 (0.0895) | 94.2076 (0.0712) | 94.5859 (0.1502) | −0.13% | +0.27% |
| Mean Weber contrast | 0.1791 (0.0011) | 0.1776 (0.0009) | 0.1823 (0.0019) | −0.84% | +1.80% |
| Michelson contrast within target | 0.5691 (0.0184) | 0.5496 (0.0647) | 0.6022 (0.0761) | −3.42% | +5.82% |

**Alt text:** The table lists mean greyscale intensity, mean Weber contrast, and Michelson contrast for Noise, Letters, and Faces, with SDs in parentheses and percentage differences for Letters and Faces relative to Noise.

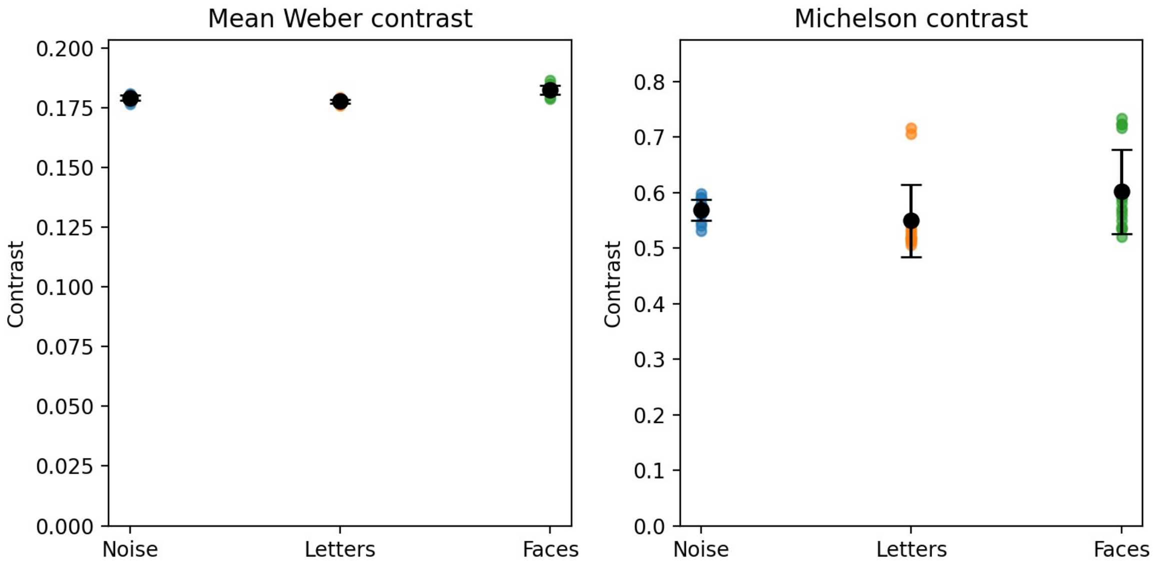


**Figure S1.1.** Per-image values of the two contrast measures. Each point is one stimulus image; the black symbol gives the category mean with ±1 SD across the 16 images of that set. Both vertical axes start at zero, so that the small differences between the category means are not visually magnified; the corresponding values are given in Table S1.1. Both measures rank the categories faces > noise > letters, whereas the observed disappearance rate ordering was faces > letters > noise.

**Alt text:** Two side-by-side plots show contrast values for Noise, Letters, and Faces. The left plot is titled “Mean Weber contrast” and has individual coloured points near 0.18 with black mean-and-error markers. The right plot is titled “Michelson contrast” and shows a wider vertical spread of points, especially for Letters and Faces, with black mean-and-error markers.

### 3. Exemplar correspondence between the letter and noise sets

Each letter was superimposed on one of the same 16 phase-randomized backgrounds that constitute the noise condition, though not in file name order. The correspondence is recovered by correlating each letter image with each noise image inside the aperture, after standardizing the pixel values within each image and then trimming the 25% most discrepant pixels, so that the letter stroke itself cannot drive the match. This one analysis uses the native 420 × 420 files rather than the presented images, because the finer sampling separates the candidate pairings more sharply. The recovery is unambiguous: the correct partner gives $r$ between 0.923 and 0.998, the best competing pairing never exceeds $r = 0.644$, the smallest margin over the runner-up is 0.33, and the recovered mapping is a bijection. It is listed in Table S1.2. Repeating the same procedure on the presented 163 × 163 images returns the identical mapping, with slightly different correlation values, so the correspondence itself does not depend on the resolution at which it is recovered.

This has a useful consequence. The letter set is the noise set with a letter superimposed and the whole image renormalized, so the letter versus noise comparison can be made within-background: for each background it is possible to measure exactly what adding the letter did. Over the 16 true partners, superimposition and renormalization lowered mean Weber contrast by 0.84% ($t(15) = -8.44$, $p < .001$) and left Michelson contrast within the target statistically indistinguishable (−3.42%, $t(15) = -1.40$, $p = .18$). The mean Weber difference is systematic, and it is small.

The face versus noise correspondence cannot be recovered with the same confidence. Phase randomization preserves the amplitude spectrum, so the source face of each noise image should in principle be identifiable from it, but circular cropping and renormalization degrade the spectrum enough that the candidate partners are separated by only a few thousandths of a correlation unit. Because the individual assignments are not secure, we do not report a face-to-noise exemplar mapping at all. The face set is compared with the noise and letter sets descriptively, without paired inference, and no claim in the main text depends on such a mapping.

**Table S1.2.** Recovered exemplar correspondence between the letter set and the phase-randomized backgrounds that constitute the noise set, with the correlation of the correct partner and of the best competing pairing.

| Letter | Background | r | Next best r | Letter | Background | r | Next best r |
|---|---|---|---|---|---|---|---|
| letter_01 | noise_02 | 0.938 | 0.526 | letter_09 | noise_10 | 0.979 | 0.568 |
| letter_02 | noise_04 | 0.966 | 0.557 | letter_10 | noise_11 | 0.989 | 0.530 |
| letter_03 | noise_05 | 0.958 | 0.555 | letter_11 | noise_12 | 0.957 | 0.606 |
| letter_04 | noise_06 | 0.923 | 0.597 | letter_12 | noise_13 | 0.996 | 0.560 |
| letter_05 | noise_07 | 0.998 | 0.620 | letter_13 | noise_14 | 0.997 | 0.597 |
| letter_06 | noise_08 | 0.997 | 0.610 | letter_14 | noise_15 | 0.995 | 0.644 |
| letter_07 | noise_01 | 0.995 | 0.619 | letter_15 | noise_16 | 0.986 | 0.596 |
| letter_08 | noise_09 | 0.998 | 0.575 | letter_16 | noise_03 | 0.988 | 0.569 |

**Alt text:** The table lists the recovered one-to-one correspondence between 16 letter images and 16 phase-randomized noise backgrounds. For each pairing it gives the correlation with the matched background and the correlation of the next-best alternative.

### 4. How large a contrast difference would be needed?

Bonneh et al. (2014) measured the dependence of MIB on target contrast parametrically, using a Gaussian luminance target in the upper-left quadrant and a rotating grid of cross-shaped mask elements, a configuration comparable to the present one. They reported that the rate of disappearance events doubles across a tenfold increase in target contrast, from 8% to 80%. Treating this as an approximate power law relation gives an exponent of log(2)/log(10) = 0.301. Table S1.3 applies this relation in both directions. It is used as an order-of-magnitude benchmark and not as a formal bound, because Bonneh and colleagues used a photometrically calibrated display and a simple Gaussian target rather than a textured image.

**Table S1.3.** Contrast difference required to produce the observed effects under the Bonneh et al. (2014) contrast function, compared with the differences actually measured, and the rate change predicted from those measured differences.

| | Faces vs noise | Letters vs noise |
|---|---|---|
| Observed disappearance rate ratio | 1.090 | 1.044 |
| Contrast difference this would require | +33.1% | +15.5% |
| Measured difference, mean Weber contrast | +1.80% | −0.84% |
| Measured difference, Michelson contrast within target | +5.82% | −3.42% |
| Predicted rate change, mean Weber contrast | +0.54% | −0.25% |
| Predicted rate change, Michelson contrast within target | +1.72% | −1.04% |

**Alt text:** The table compares Faces versus Noise and Letters versus Noise for the observed disappearance-rate ratio, the contrast difference required by the Bonneh et al. benchmark, the measured Weber and Michelson contrast differences, and the rate changes predicted from those measured differences.

On the measure that the MIB literature has actually manipulated, the residual difference between faces and noise corresponds to about 6% of the observed effect for faces; on Michelson contrast, which separates the categories more, it corresponds to about 19%. Neither measure, however, reproduces the observed ordering. Both rank the categories faces > noise > letters, and therefore both predict a lower disappearance rate for letters than for noise, whereas letters in fact disappeared more often than noise. This conclusion does not depend on the resampling rule: across the four alternative interpolation rules used as a check, the predicted rate change never exceeded 1.8% in magnitude for either category, against observed effects of 9.0% for faces and 4.4% for letters, and in every case the prediction for letters carried the wrong sign. This is the same conclusion that Figure S1.1 shows graphically.

### 5. Spatial frequency content

The noise images were generated by randomizing the phase spectrum of the face images while preserving their amplitude spectrum (Näsänen, 1999), so faces and noise are identical in amplitude spectrum prior to circular cropping, but not after it, since cropping retains different content from a face than from its phase-randomized counterpart. Figure S1.2a shows the rotational average of the Fourier amplitude spectrum of the contrast image, computed per image and averaged within categories (Cooley & Tukey, 1965; Field, 1987; Willenbockel et al., 2010); Figure S1.2b expresses the same data as the deviation from the noise baseline.

Averaged across spatial frequencies, the letter set deviates from the noise baseline by 4.20% and the face set by 9.52%. One descriptive reference for the scale of these differences is the standard error of the mean amplitude across the 16 noise images, which is 3.20%. The letter set therefore lies little more than one standard error from the noise baseline, whereas the face set lies about three, and the letter set is the more closely matched of the two in every one of the five frequency bands of Table S1.4. This is a descriptive benchmark and not a formal equivalence test, and it does not amount to spectral equivalence. The measured pattern is that the letter set, although artificial, lies closer to the spectrum of the noise images than the face set does.

Table S1.4 also shows that the two categories deviate from the noise baseline in opposite directions across four of the five bands, the letter set carrying more amplitude and the face set less, while both categories showed higher disappearance rates than noise. A monotonic dependence of disappearance rate on spectral amplitude therefore cannot reproduce the observed ordering.

**Table S1.4.** Deviation from the noise baseline in the rotationally averaged amplitude spectrum, by spatial frequency band (ratio of summed amplitude).

| Spatial frequency band | Letters vs noise | Faces vs noise |
|---|---|---|
| 0.5 to 2 cycles/deg | −2.03% | −13.41% |
| 2 to 5 cycles/deg | +4.58% | −8.18% |
| 5 to 10 cycles/deg | +7.76% | −11.71% |
| 10 to 15 cycles/deg | +5.14% | −12.01% |
| 15 to 40 cycles/deg | +2.80% | −8.53% |
| Mean absolute deviation, all frequencies | 4.20% | 9.52% |
| For comparison: SEM within the noise set | 3.20% | 3.20% |

**Alt text:** The table lists percentage deviations from the Noise amplitude-spectrum baseline for Letters and Faces across five spatial-frequency bands, followed by the mean absolute deviation across frequencies and the SEM within the Noise set.

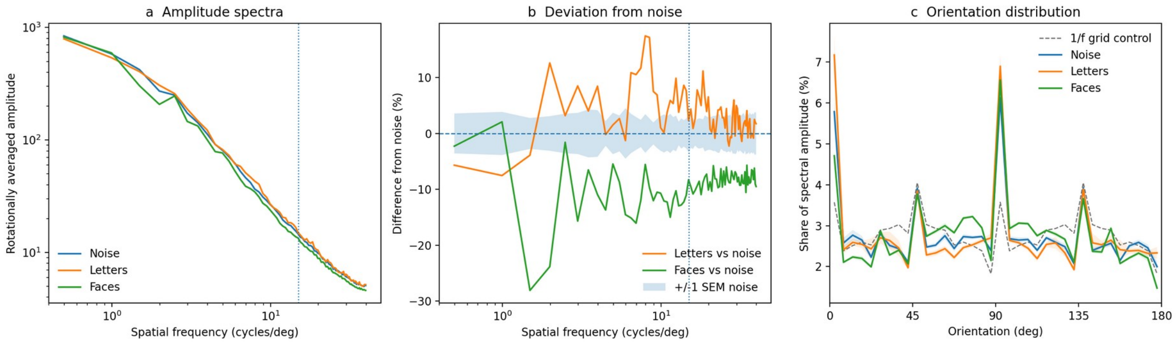


**Figure S1.2.** (a) Rotationally averaged Fourier amplitude spectra of the three categories; the dotted vertical line marks 15 cycles per degree. (b) Deviation from the noise baseline for the letter and the face set, in the category colours of (a) and (c); the shaded band gives ±1 standard error of the mean amplitude of the noise set, shown as a descriptive reference band rather than as an equivalence criterion; it is not the spread across those images, which is four times larger. (c) Distribution of Fourier amplitude across orientations, in 36 bins of 5 deg and expressed as the percentage share of the total non-DC amplitude of each image, with shaded bands giving ±1 standard error across the 16 images of each set. The grey dashed curve is the same histogram computed for an isotropic 1/f amplitude spectrum sampled on the same 163 × 163 Fourier grid. It is not flat, because a square discrete grid does not sample orientations uniformly, and it is therefore the appropriate reference against which to read the three measured curves.

**Alt text:** Three side-by-side line plots summarize image spectra. Panel (a) shows three downward-sloping amplitude-spectrum curves for Noise, Letters, and Faces on logarithmic axes, with a dotted vertical line at 15 cycles per degree. Panel (b) shows orange and green percentage-deviation curves around a horizontal zero line with a light-blue shaded reference band. Panel (c) shows coloured orientation-distribution curves and a grey dashed reference curve, with prominent peaks near the cardinal orientations.

The dotted line in Figure S1.2 marks 15 cycles per degree, which is used here only as a descriptive cutoff. Restricting attention to the bands below that line does not change the overall descriptive pattern: across those four bands the letter set departs from the noise baseline by at most 7.76%, against a noise standard error of 3.20%, whereas the face set departs by between 8.18% and 13.41%. The two categories deviate in opposite directions in three of these four bands; in the 0.5–2 cycles/deg band both fall below the noise baseline. We do not treat 15 cycles per degree as a physiological or resolution limit; the restriction only shows that the same broad spectral differences are present in the lower-frequency range, and it is not evidence that the target categories were discriminable in the periphery, which was assessed behaviourally in Experiment A of the main text.

The rotational average does not capture orientation. Figure S1.2c gives the distribution of Fourier amplitude across orientations (Hansen & Essock, 2004), computed on the same presented 163 × 163 contrast images in 36 bins of 5 deg, excluding the DC component and normalized to sum to 100% within each image. The cardinal-to-oblique ratio, defined as the amplitude share in the bins whose centres lie within 15 deg of 0 and 90 deg divided by the share in the bins whose centres lie within 15 deg of 45 and 135 deg, was computed for each image and then averaged within categories. It is 1.180 (SD = 0.139) for the noise images, 1.268 (SD = 0.246) for the letters and 1.121 (SD = 0.061) for the faces. The same ratio computed for an isotropic 1/f amplitude spectrum sampled on the same discrete grid is 0.809, not 1, because a square Fourier grid does not sample orientations uniformly; the raw values should therefore not be read as percentages above a flat orientation histogram, and only the differences between categories are interpreted here. Relative to the noise baseline the letter set is about 7% more anisotropic and the face set about 5% less, the letter value lying in the direction that the predominantly vertical and horizontal strokes of Roman characters imply. The spread across exemplars is large relative to these category differences, and the orientation summary is accordingly treated as descriptive; no inferential claim depends on it. The residual anisotropy of the letter set is noted in the Limitations of the main text.

### 6. References

All works cited in this supplement are listed in the consolidated reference list of the main manuscript.
