## Supplementary_Material_2 for "Stimulus category is associated with conscious access during motion-induced blindness"

Descriptive summaries and robustness analyses

### 1. Descriptive summaries by stimulus category

All values are based on the 1,632 experimental trials of Experiment B (34 participants × 48 trials); the single practice trial of each participant was excluded. Disappearance rate is computed per participant as the total number of disappearance onsets divided by the total visible time, and then averaged across participants. Episode durations are computed over completed episodes only.

**Table S2.1.** Trial- and episode-level descriptives by stimulus category.

| Measure | Noise | Letters | Faces |
|---|---|---|---|
| Disappearance rate (per s visible), M (SEM) | 0.149 (0.016) | 0.154 (0.016) | 0.161 (0.016) |
| Disappearance rate (per s of trial), M (SEM) | 0.122 (0.011) | 0.127 (0.011) | 0.132 (0.011) |
| Invisibility fraction, M | 0.132 | 0.134 | 0.135 |
| Invisible episode duration (s), M | 1.044 | 1.043 | 0.980 |
| Invisible episode duration (s), Mdn | 0.763 | 0.745 | 0.746 |
| Completed invisible episodes, n | 2,911 | 3,033 | 3,146 |
| Visible episode duration (s), M | 5.329 | 5.166 | 5.086 |
| Visible episode duration (s), Mdn | 3.613 | 3.470 | 3.492 |
| Completed visible episodes, n | 2,984 | 3,106 | 3,240 |

**Alt text:** The table summarizes disappearance rates, invisibility fraction, invisible episode durations, visible episode durations, and completed episode counts for Noise, Letters, and Faces. Means, medians, and SEMs are provided where applicable.

**Table S2.2.** Mean invisibility fraction across the four experimental blocks.

| Block | 1 | 2 | 3 | 4 |
|---|---|---|---|---|
| Mean invisibility fraction | 0.174 | 0.125 | 0.109 | 0.126 |

**Alt text:** The table lists mean invisibility fraction for experimental Blocks 1 through 4, with one value shown for each successive block.

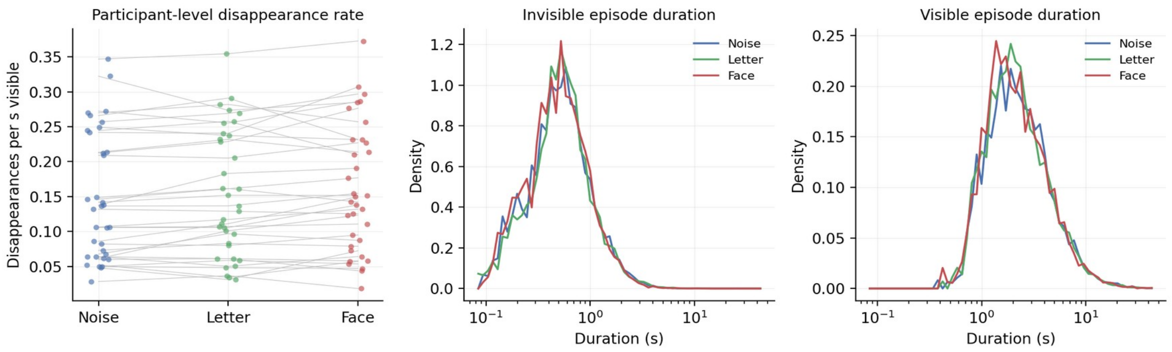


**Figure S2.1.** Raw distributions by stimulus category. (a) Participant-level disappearance rate, with each line connecting the three category means of one participant. (b) Distribution of completed invisible episode durations, plotted on a logarithmic time axis. (c) Distribution of completed visible episode durations, on the same axis. The distributions are strongly skewed and overlap substantially, which is why the inferential analyses are carried out on log-transformed durations.

**Alt text:** Three side-by-side plots. Panel (a) shows participant-level disappearance rates for Noise, Letter, and Face, with small points linked by faint lines across the three categories. Panel (b) shows overlapping density curves for invisible-episode durations on a logarithmic horizontal axis. Panel (c) shows overlapping density curves for visible-episode durations on the same type of logarithmic axis.

### 2. Robustness analyses

The following sensitivity analyses were computed to establish whether the reported conclusions depend on specific analytic choices. Primary model estimates are shown alongside alternative exposure definitions, nuisance controls, boundary handling and participant-clustered marginal refits. All models use noise as the reference category. Visible episode duration is an exploratory outcome. Its model carries the same participant random intercept and the same fixed effects of stimulus category and centred block as the invisible episode model, but no covariate for the duration of the preceding episode; it is labelled accordingly below. The alternatives correspond to established model classes: Poisson rate models with exposure offsets (Frome, 1983), marginal generalized estimating equations for clustered repeated observations (Liang & Zeger, 1986), random-intercept mixed models for repeated and unbalanced observations (Laird & Ware, 1982; Pinheiro & Bates, 2000), and fractional response quasi-likelihood for outcomes bounded between 0 and 1 (Papke & Wooldridge, 1996).

**Table S2.3.** Robustness analyses for the effect of stimulus category, with noise as the reference category.

| Analysis | Letters vs noise | Faces vs noise |
|---|---|---|
| Disappearance rate, Poisson GEE with log(visible time) offset (primary) | IRR = 1.044 [1.004, 1.087], p = .031 | IRR = 1.090 [1.027, 1.157], p = .005 |
| Disappearance rate, count per full 45 s trial | IRR = 1.041 [1.007, 1.076], p = .019 | IRR = 1.086 [1.032, 1.143], p = .002 |
| Disappearance rate with rotation and shift direction nuisance covariates | IRR = 1.042, p = .037 | IRR = 1.089, p = .005 |
| Disappearance rate after excluding 6 trials with 8 to 14 ms end time overshoot | IRR = 1.048 [1.008, 1.090], p = .018 | IRR = 1.088 [1.024, 1.156], p = .006 |
| Invisible duration, mixed-effects model (primary) | β = −0.001 [−0.032, 0.031], p = .974 | β = −0.046 [−0.078, −0.015], p = .004 |
| Invisible duration, mixed model without preceding visible covariate | β = 0.001 [−0.031, 0.033], p = .943 | β = −0.044 [−0.075, −0.012], p = .007 |
| Invisible duration, participant-clustered Gaussian GEE (exchangeable) | β = −0.001 [−0.026, 0.025], p = .969 | β = −0.047 [−0.084, −0.009], p = .016 |
| Visible duration, mixed-effects model (exploratory) | β = −0.033 [−0.068, 0.001], p = .059 | β = −0.053 [−0.087, −0.018], p = .003 |
| Visible duration, participant-clustered Gaussian GEE (exchangeable) | β = −0.033 [−0.066, −0.001], p = .046 | β = −0.053 [−0.096, −0.010], p = .016 |
| Invisibility fraction, logit mixed-effects model (primary) | β = 0.225 [−0.107, 0.558], p = .184 | β = 0.229 [−0.104, 0.561], p = .178 |
| Invisibility fraction, fractional response GEE (exchangeable, no clipping) | β = 0.020 [−0.038, 0.079], p = .495 | β = 0.030 [−0.053, 0.113], p = .482 |

**Alt text:** The table lists primary and robustness-model estimates for Letters versus Noise and Faces versus Noise across disappearance rate, invisible duration, visible duration, and invisibility fraction. Each entry reports the corresponding effect estimate, confidence interval where available, and p value.

The originally planned repeated-measures analysis of variance, computed on participant-level disappearance rates within the classical repeated-measures and profile data framework (Greenhouse & Geisser, 1959), also yielded a reliable effect of stimulus category, $F(2, 66) = 4.286$, $p = .0178$, converging with the primary Poisson GEE result.

The exploratory direct comparison between the face and letter categories, obtained by re-levelling the same primary models, gave IRR = 1.043, 95% CI [0.987, 1.103], $p = .132$ for disappearance rate and $\beta = -0.046$, 95% CI [−0.077, −0.015], $p = .004$ for invisible episode duration. Faces and letters therefore did not differ reliably in disappearance rate, whereas invisible episodes were shorter for faces. These comparisons are exploratory, because the planned contrasts used noise as the common reference.

**Raw event reconstruction.** Event-level analyses were reconstructed from the DisappearTimes and ReappearTimes event fields, not from the legacy derived DisappearanceDurations column in the raw CSV files. When a trial ended with an unmatched disappearance, that legacy derived field can contain a negative bookkeeping value, which is not a negative perceptual duration. The analysis instead marks an unmatched terminal disappearance as right-censored at trial end, following standard survival analysis handling of incomplete event times (Cox, 1972; Kleinbaum & Klein, 2012), as described in the Methods. Censoring is handled inferentially only in the Cox model of latency to the first disappearance. The primary log duration mixed model does not use a survival likelihood and is therefore fitted to completed episodes only: of the 9,330 disappearance onsets, 9,090 were followed by a reappearance within the trial and enter the model, while the remaining 240 were still ongoing when the trial ended and are excluded.

**Model fitting.** The primary invisibility fraction analysis clips IF to $[1 \times 10^{-6}, 1 - 10^{-6}]$, applies a logit transform, and fits a participant random-intercept mixed model. Because 150 of 1,632 trials have IF = 0, the fractional response GEE in Table S2.3 is reported as a boundary-safe sensitivity check rather than as a replacement primary analysis; fractional response quasi-likelihood is designed to accommodate proportions at the boundaries without arbitrary displacement (Papke & Wooldridge, 1996), while GEE accounts for clustering across repeated trials (Liang & Zeger, 1986). For the duration mixed models, converged BFGS solutions gave participant random intercept variances of 0.132 (invisible) and 0.175 (visible), and conjugate gradient and Powell optimization reproduced the same substantive solutions. These optimizer checks concern numerical convergence of the same likelihood rather than alternative statistical models; BFGS and conjugate gradient methods are standard numerical optimization families (Nocedal & Wright, 2006), while the derivative-free Powell search is the method introduced by Powell (1964). The Cox proportional hazards model for latency to the first disappearance is fitted at the trial level with standard errors clustered by participant (Lin & Wei, 1989); it is reported in the main text rather than among the refits above, because the alternative specifications listed there do not apply to a time-to-event outcome.

**Multiplicity.** The sequentially rejective Holm correction (Holm, 1979) was applied separately to the two planned noise baseline contrasts within each outcome family. Adjusted p values (letters, faces) were: disappearance rate (.031, .010), invisible duration (.974, .007), visible duration (.059, .005), and invisibility fraction (.355, .355). Both disappearance rate contrasts and the face duration contrasts remain reliable, while no primary null result becomes reliable after correction.

**Interpretation of the letter contrast for visible duration.** This contrast is model-sensitive, with $p = .059$ in the primary mixed-effects model and $p = .046$ in the exchangeable GEE sensitivity refit. It is therefore not treated as a stable finding. The face versus noise effects on both invisible and visible duration are supported in both specifications.

### 3. References

All works cited in this supplement are listed in the consolidated reference list of the main manuscript.
