## Supplementary_Material_3 for "Stimulus category is associated with conscious access during motion-induced blindness"

An independent eye-tracking control dataset

### 1. Scope

In Experiment B the eye tracker was used for online fixation control only. Gaze was shown continuously on the experimenter's control monitor and participants were re-instructed and re-calibrated when drift was observed, but no samples were written to disk. There is therefore no gaze record of Experiment B to analyse, and none is reported here.

This supplement reports instead an independent dataset from the same laboratory in which gaze was recorded throughout. It comes from a motion-induced blindness (MIB) experiment run in the same room, on the same display at the same viewing distance, with the same EyeLink 1000 Plus unit, and with the target centre rendered at the same pixel eccentricity as in Experiment B. Three of its five target classes correspond to those of the main study: phase-randomized control images, faces, and Roman letters.

Two constraints apply throughout: these are not the eye movements of Experiment B, and nothing reported here is a replication of Experiment B. Section 6 states the conclusion the dataset supports and its four limits.

This supplement is a control analysis and not an additional experiment of the present study. The data were recorded earlier, for a broader purpose that also included inverted faces, letter-like glyphs and four target locations (Section 2.1), and the behavioural design could have limited both sensitivity to the category question and comparability with Experiment B (Section 2.2). The dataset is therefore reported for the one question it can address, namely whether the three target classes of Experiment B show reliable differences in how they are looked at, on the measures recorded here and under the viewing conditions of this experiment, at this eccentricity and within an inducer field of nearly the same size, and not as independent evidence for or against the effects reported in the main text. It is nevertheless reported at full length, with the linkage and quality-control audit, the three fixation-window definitions, the effect sizes and their bootstrap intervals, and the archived analysis package of Section 7, so that a control offered in place of a missing gaze record can be checked in detail.

2. The control experiment

**2.1 What it was for, and what it delivered.** The data were collected between 23 May and 21 June 2024 on fourteen testing days, in a study designed around a broader question than the present one. Target class had five levels (phase-randomized control images, upright faces, the same faces inverted, Roman letters, and unfamiliar, letter-like glyphs), chosen to separate a face-specific effect from a more general effect of familiarity or of figural structures. Gaze was recorded on every trial.

Analysed at the participant level across all five classes, target class had no reliable effect on any measure of MIB dynamics (Table S3.7): response-enabled invisibility fraction, $F(4, 136) = 1.94$, $p = .105$; disappearance onset rate per second of response-enabled visible time, $F(4, 136) = 0.57$, $p = .687$; disappearance reports per trial, $F(4, 136) = 0.40$, $p = .812$. None of the ten pairwise contrasts survived Holm correction for any measure in Table S3.7; the smallest adjusted value was $p = .060$, for control versus letters on total invisible duration.

2.2 Design features that may have limited sensitivity. The trial lasted 30 s rather than 45 s, so that with sixteen trials per class the planned exposure per class was 480 s rather than 720 s. The target appeared in one of four oblique quadrants rather than always in the upper left, which adds between-location variance to a within-participant contrast. And the target was rendered within a 108 pixel frame (1.33 deg), with a visible oval aperture narrower still (Section 3.3), rather than the 163 pixel circular disc (2.01 deg) used in Experiment B, and its peripheral discriminability was not separately assessed in a discrimination task of the kind reported as Experiment A. Behavioural values remained descriptively lower after recomputing rate and invisibility fraction over the response-enabled part of the trial: mean invisibility fraction ranged from 0.103 to 0.113 against 0.132 to 0.135 in Experiment B, disappearance onset rate ranged from 0.136 to 0.143 per second of response-enabled visible time against 0.149 to 0.161, and 12.0% of trials contained no disappearance report at all.

2.3 Why it is appropriate as a control dataset. The same sessions produced 1000 Hz binocular gaze, blink, saccade and pupil records for control, face and letter targets, at the same eccentricity, in the same room, on the same tracker, and within an inducer field of nearly the same size (Table S3.1). Two further properties matter for the use made of the data here. Because the behavioural outcome was null, the eye records cannot have been selected in favour of any behavioural pattern. And because the target occupied four locations rather than one, the analysis tests whether any category-related gaze bias generalizes across quadrants rather than only at a single fixed target location; a bias confined to one location could correspondingly be diluted.

**Table S3.1.** Experiment B and the control dataset compared. Visual angles are given at 81.00 pixels per degree throughout (Section 3.2). Trials per class and exposure per class are planned quantities; in the control dataset four participants contributed fewer than four blocks and therefore less than the planned exposure (Section 3.1).

| Feature | Experiment B (main text) | Control dataset (this supplement) |
|---|---|---|
| Participants | 34 | 35 (31 with all four blocks) |
| Target classes | 3: noise, letters, faces | 5: control, faces, inverted faces, letters, glyphs |
| Trials per class per participant (planned) | 16 | 16 |
| Trial duration | 45 s | 30 s |
| Exposure per class (planned) | 720 s | 480 s |
| Target location | Upper-left quadrant only | Four oblique quadrants |
| Target centre eccentricity | 583 px (7.20 deg) | 583 px (7.20 deg) |
| Target size | 163 px (2.01 deg), circular | 108 × 108 px frame (1.33 deg); visible oval 80 × 108 px (0.99 × 1.33 deg) |
| Inducer field diameter | 1368 px (16.9 deg) | 1306 px (16.1 deg) |
| Inducer element size | 1.0 deg crosses | 0.67 deg crosses |
| Inducer rotation | 60 deg/s | 133 deg/s |
| Background | 8-bit 80 | 8-bit 100 |
| Display and viewing distance | 2560 × 1440, 59.6 cm wide, 105.3 cm | Identical |
| Presentation software | Python 3.9.21, Pygame 2.6.1 | PsychoPy 2021.1.4, Python 3.6 |
| Eye tracking | EyeLink 1000 Plus, online control only, no samples stored | Same unit, 1000 Hz binocular, recorded to disk |
| Mean invisibility fraction | 0.132–0.135 | 0.103–0.113 (response-enabled) |
| Disappearance onset rate | 0.149–0.161 /s visible | 0.136–0.143 /s response-enabled visible |

**Alt text:** The table compares Experiment B with the independent control dataset across participants, target classes, trial structure, target and inducer geometry, display settings, presentation software, eye-tracking acquisition, and behavioural summary measures.

### 3. Method of the control experiment

**3.1 Participants.** Thirty-five participants contributed data. Mean age was 22.54 years (SD = 2.48, range 19 to 27 years); 28 were female and 7 male. Four experimental blocks were recorded and analysable for thirty-one participants, three blocks for three participants and a single block for one. Of the six blocks missing relative to four per participant, three were run but had no EyeLink file written (Section 5.1), and three were not run at all. Recruitment was conducted through the internal system of the Institute of Psychology, Friedrich Schiller University Jena. All participants had normal or corrected-to-normal vision and reported no neurological or psychiatric history. Participation was rewarded with partial course credit. Ethical approval was granted by the local ethics committee (ethics number: FSV 21/073), and all procedures were carried out in accordance with the Declaration of Helsinki. Written informed consent was obtained from each participant prior to testing.

**3.2 Apparatus, display and the two conversion factors.** Stimuli were presented on a 2560 × 1440 pixel display of 59.6 cm active width at a viewing distance of 105.3 cm, the same configuration as in Experiment B, with the window opened in PsychoPy height units at a nominal refresh rate of 165 Hz.

Visual angles are reported here at 81.00 pixels per degree, the factor used throughout the main text, which the presentation scripts of both experiments derive from the full screen width. The offline gaze analysis used a different factor: the per-trial resolution value the EyeLink writes to the END record where it was available, and otherwise 78.95 pixels per degree, which is the tangent-based value at the centre of the screen and matches the resolution the tracker itself reported. The two differ by 2.6%, because the first is an average across the width of a flat display and the second is the local value at its centre. The difference is confined to the fixation-window criteria, which are stated in the analysis units: the 1.5, 2 and 3 deg windows correspond to 118.4, 157.9 and 236.9 pixels, that is, to 1.46, 1.95 and 2.92 deg in the units of the main text. No conclusion in Section 5 depends on which factor is adopted.

**3.3 Target images.** The seventeen target images were 400 × 400 pixel RGBA PNGs, greyscale in content (R = G = B at every pixel), sharing a single binary alpha mask. The mask is an upright oval rather than a circle: measured on the archived files it spans the full 400 pixel height of the frame and 296 pixels of its width, truncated by the frame at top and bottom, and leaves 94,492 pixels opaque, 59.06% of the frame, against the 78.5% that an inscribed circle would occupy; all seventeen images carry the identical mask. Drawn at 108 × 108 pixels (Section 3.4), the visible target therefore measured 80 × 108 pixels, 0.99 × 1.33 deg, where the target of Experiment B was a circular disc of 163 pixels, 2.01 deg. The set comprised one phase-randomized control image, four faces, those four faces inverted, four letters and four glyphs, all of which had passed a SHINE-based intensity and contrast equalization (Willenbockel et al., 2010). Figure S3.1 shows one exemplar of each class (faces: synthetic illustration).

The five classes differ in how the visible form was produced. The control image is a phase-randomized image; the letter and glyph images carry a dark figure drawn over a phase-randomized background of the same kind, so that they share the background type of the control class rather than its amplitude spectrum, since drawing a figure over that background alters the spectrum; the size of such differences is quantified for the stimulus set of the main experiment in Supplementary Material S1, Section 5; the face images are photographic, and the inverted faces are the same four photographs rotated by 180 deg. Every value reported below was measured on the final PNG files as they were uploaded to the display, inside the aperture and after equalization, so that the matching is verified on the presented images rather than assumed from the generation procedure.

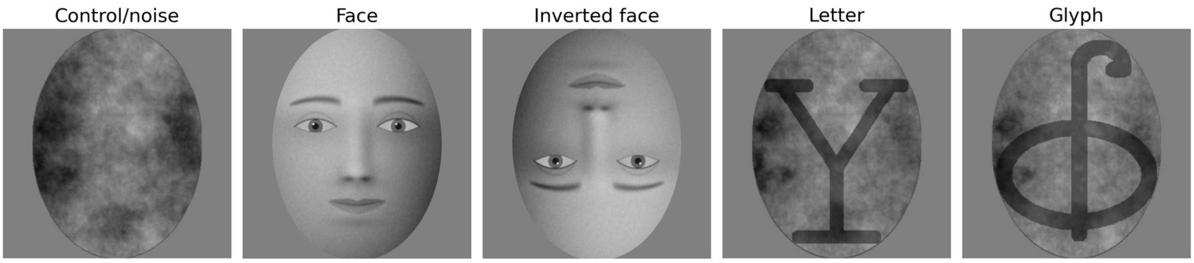


**Figure S3.1.** *Target classes of the control dataset.* One exemplar of each of the five classes at the common oval aperture. **Note:** the Face and Inverted face exemplars show a synthetic, computer-generated face created for illustration only, which does not depict a real person; the actual face stimuli were greyscale photographs and are not reproduced here in order to protect the identity of the depicted persons. The control, letter and glyph exemplars are shown as stored. Control images are phase-randomized; letters and glyphs are drawn over a phase-randomized background of the same kind; inverted faces are the upright faces rotated by 180 deg. All seventeen images were equalized for mean intensity and contrast and measured after equalization (Table S3.2). The images are reproduced at a common scale and not at their presented size of 108 × 108 pixels.

**Alt text:** Five greyscale oval target examples are arranged in one row and labelled Control/noise, Face, Inverted face, Letter, and Glyph. The control is a textured noise pattern; the face is a synthetic, computer-generated face (shown in place of the photographic stimuli), upright and upside down in the next two panels; the letter is a dark Y over a textured background; and the glyph is a dark unfamiliar symbol over a textured background.

Table S3.2 gives the measurements for all seventeen images. The mean 8-bit code value across the set was 101.496, with a total range of 0.051 code values from the darkest to the brightest image and a coefficient of variation of 0.017%; the class means were 101.501 for the control image, 101.482 for faces and 101.519 for letters. As in Supplementary Material S1, mean Weber contrast relative to the display background was taken as the primary scalar contrast measure, defined as $(\mu - B)/B$ with $B = 100$; it was 0.01501 for the control image, 0.01482 on average for faces, and 0.01519 for letters. RMS contrast, defined as the population standard deviation divided by the image mean, and Michelson contrast within the aperture were retained as complementary measures; RMS contrast averaged 28.48% and spanned 28.29% to 29.20%, while Michelson contrast spanned 0.848 to 0.856. Within the control, face and letter images the largest deviation from the control image in RMS contrast was 0.74% in relative terms, and for the letters specifically at most 0.39%. A single glyph image lay outside that band at +2.44%; glyphs enter the omnibus tests across all five classes in Section 5 but not the critical pairwise contrasts of Table S3.6, which are confined to the control, face and letter classes. The three classes that carry the critical contrasts are therefore closely matched in mean intensity and in mean Weber contrast relative to the display background, with the complementary RMS and Michelson measures likewise showing only small differences.

The background against which the targets were drawn was 8-bit 100, so that the mean target code values were close to the background code value. As in Supplementary Material S1, these are code values and not photometrically measured luminances. At presentation, each image was uploaded as an OpenGL texture and drawn at 108 × 108 pixels with bilinear interpolation.

**Table S3.2.** Intensity and contrast of the seventeen target images of the control dataset, measured inside the shared oval aperture on the final PNG files. Code values are 8-bit greyscale. RMS contrast is the population standard deviation of the code values divided by their mean; Michelson contrast is computed within the aperture. The last column gives each image's RMS contrast as a relative deviation from the control image. The background against which the targets were drawn was code value 100.

| Class | Exemplar | Mean code value | RMS contrast (%) | Michelson contrast | Deviation from control in RMS contrast (%) |
|---|---|---|---|---|---|
| Control | — | 101.501 | 28.505 | 0.856 | 0.00 |
| Faces | f1 | 101.472 | 28.295 | 0.856 | −0.74 |
| | f3 | 101.482 | 28.311 | 0.856 | −0.68 |
| | m3 | 101.488 | 28.368 | 0.856 | −0.48 |
| | m4 | 101.487 | 28.341 | 0.856 | −0.57 |
| Letters | d | 101.523 | 28.610 | 0.856 | +0.37 |
| | f | 101.515 | 28.479 | 0.856 | −0.09 |

| Class | Exemplar | Mean code value | RMS contrast (%) | Michelson contrast | Deviation from control in RMS contrast (%) |
|---|---|---|---|---|---|
| | p | 101.518 | 28.617 | 0.856 | +0.39 |
| | y | 101.522 | 28.473 | 0.856 | −0.11 |
| Inverted faces | f1 | 101.473 | 28.293 | 0.856 | −0.74 |
| | f3 | 101.479 | 28.314 | 0.856 | −0.67 |
| | m3 | 101.491 | 28.366 | 0.856 | −0.49 |
| | m4 | 101.483 | 28.346 | 0.856 | −0.56 |
| Glyphs | d | 101.512 | 28.580 | 0.856 | +0.26 |
| | f | 101.504 | 28.483 | 0.856 | −0.08 |
| | p | 101.476 | 29.200 | 0.848 | +2.44 |
| | y | 101.510 | 28.515 | 0.856 | +0.04 |
| All 17 images | mean | 101.496 | 28.476 | 0.856 | — |

**Alt text:** The table lists all 17 control-dataset images by class and exemplar, with mean 8-bit code value, RMS contrast, Michelson contrast, and relative RMS-contrast deviation from the control image.

3.4 MIB inducer, design and trial structure. The inducer was a texture of 141 blue cross-shaped elements arranged within a circular field, drawn at 3600 × 3600 pixels and therefore extending beyond the display on every side. The source file is anisotropically pre-distorted, and the scaling applied to it (2.691 horizontally, 3.406 vertically) restores the field to a circle on screen: measured on the rendered image the field spans 1308 pixels horizontally and 1304 vertically, a diameter of 16.1 deg against 16.9 deg in Experiment B. Individual elements measure 0.67 deg against 1.0 deg. The field was rotated by cycling through 360 one-degree orientation steps every 2.7 s, one full revolution at 133 deg/s against 60 deg/s in Experiment B, with the direction randomized on each trial.

The target centre lay 583 pixels from fixation, 7.20 deg, the same distance as in Experiment B, on one of the four oblique diagonals; the polar angle was drawn uniformly from five integer values spanning ±2 deg around the diagonal. Taking the 54 pixel half-height of the target frame as a bounding radius, the outer edge of the target reached 637 pixels, 7.86 deg, which is 16 pixels inside the outer boundary of the inducer field at 653 pixels, 8.06 deg; because the visible aperture is an oval narrower than its frame (Section 3.3), the clearance along the oblique meridian is in fact larger than this. The corresponding margin in Experiment B was 0.24 deg. Fixation was marked by a cyan spot 14.4 pixels (0.18 deg) across that blinked at 1 Hz.

Each block presented the twenty combinations of five target classes and four quadrants in a fixed order, drawing a different exemplar of each class in each block; four blocks gave 80 trials, sixteen per class. A trial began with a fixation interval drawn uniformly from 1.5 to 2.0 s and continued with a 30 s stimulus interval. Participants pressed Z at a reported disappearance and M at a reported reappearance; responses did not alter the display.

**3.5 Eye-tracking acquisition and online fixation control.** Recording used an EyeLink 1000 Plus (EyeLink II CL v5.50, 16 June 2022; camera EyeLink GL v1.2, sensor AG7; serial CLG-BIE17) in binocular mode at 1000 Hz. All 134 recorded blocks came from this one unit. The tracker wrote LEFT, RIGHT, FIXATION, SACCADE, BLINK, MESSAGE, BUTTON and INPUT events and LEFT, RIGHT, GAZE, HREF, RAW, AREA, HTARGET, GAZERES, BUTTON, STATUS and INPUT sample fields. A nine-point calibration (HV9) was run at the start of each block, and re-calibration and validation were available at every block boundary. Recording was started and stopped on each trial, and a separate EDF file was written for each block.

Two gaze-contingent regions were defined at the start of every trial. The first was centred on fixation and gated the response period: key presses were not recorded until gaze had remained inside it continuously for 1.5 s, after which the gate stayed open for the remainder of the trial. Because the script converts screen pixels into an internal coordinate space in which the display is treated as 1.6 units wide and 1.0 units high, whereas its true aspect ratio is 1.778, this nominally circular region is in fact an ellipse on screen, with semi-axes of 240 pixels horizontally and 216 pixels vertically, that is, 2.96 deg by 2.67 deg. The second region was centred on the target; it was updated on every frame, but the branch that would have acted on it is commented out in the production script, so it neither aborted nor flagged any trial.

Two consequences follow, and both matter for the interpretation of Section 5. No trial was rejected online on the basis of gaze, and no automatic drift correction was performed between trials. Every trial in which the participant satisfied the initial fixation gate therefore contributes its full 30 s of samples, whatever the gaze did afterwards, and all quality control reported below is applied offline.

### 4. Offline analysis

**4.1 EDF conversion.** All 134 EDF files were converted to ASCII with SR Research EDF2ASC 4.2.1197.0 using edf2asc -t -miss NaN -y -p <directory> <file>, which writes tab-delimited output and represents missing gaze coordinates as NaN. All 134 files converted and parsed without error; the console logs are archived. The intermediate ASCII files are not archived, because they are large and exactly reproducible from the EDF files and this command.

**4.2 Trial reconstruction and behavioural linkage.** Trials were identified from the TRIALID messages and the corresponding START and END records. The tracker-side class label was read from the !V TRIAL_VAR condition message, reported disappearances and reappearances from the dis_key_pressed and reap_key_pressed messages, and blinks and saccades from the EBLINK and ESACC records. Each eye-tracking trial was matched to a row of the PsychoPy log by participant, zero-based block index and Trial_ID, from which the target class, quadrant, response counts and response times, summed invisible duration, image file and frame rate were taken. This produced 2,680 linked trials, the full 134 × 20.

**4.3 Definition of the analysis window.** The stimulus interval of a trial is 30 s, the value set in the parameter module that the presentation script imports, and the whole of it is analysed for the eye and oculomotor measures.

Its onset in EyeLink time was taken as the TRIALID timestamp plus the interval between the two PsychoPy start markers, imageA.started minus fix_centre.started. This estimate was available for all 2,680 trials; the offset had a median of 1605.0 ms (SD 146.9, range 1245.5 to 1894.4), consistent with the programmed 1.5 to 2.0 s fixation interval. The window ran 30 s from that onset, truncated at the END record where necessary; the realized duration was 30.000 s on every trial, giving exactly 30,000 samples per trial, so no window was in fact truncated; the EDF recording interval from TRIALID to END averaged 31.715 s.

A second and independent alignment was computed as a check, by subtracting each logged response time from the timestamp of the corresponding EyeLink response message. It was available on 2,364 trials, and the several estimates within a trial agreed to a median absolute residual of 1.96 ms. Across trials this response-based onset fell a median of 1533 ms after the marker-based onset, with a minimum of 1516 ms. That displacement is the 1.5 s fixation gate of Section 3.5, because response times are counted from the moment the gate opened rather than from image onset. The two clocks therefore agree once the gate is accounted for. The analysis window is defined by the marker-based estimate, which is anchored to image onset itself. For the behavioural rate and invisibility-fraction summaries, the response-enabled window was defined as the 30 s stimulus interval minus this response-gate offset; when no response-based alignment was available, the median offset across aligned trials was used.

**4.4 Gaze metrics.** Left and right gaze positions were averaged when both eyes were valid, and the valid eye was used when only one was; a sample with no finite coordinate in either eye was counted as invalid. Distance from fixation was computed from the screen centre at (1280, 720) and converted to degrees using the per-trial resolution written by the tracker where available and 78.95 pixels per degree otherwise. For each trial we computed the fraction of valid samples, the percentage of valid samples farther than 1.5, 2 and 3 deg from fixation, the mean and median gaze distance from fixation, and the standard deviations of horizontal and vertical gaze position.

**4.5 Blinks, saccades and pupil.** Blink and saccade rates were computed as EyeLink EBLINK and ESACC event-record rates within the analysis window, divided by its duration in seconds; no cross-eye deduplication was applied. Mean saccade amplitude was taken from the ESACC records. Pupil area was averaged across valid eyes in the same way as gaze position. Baseline-corrected pupil area was the mean over the window minus the mean over its first 500 ms. Microsaccades were not extracted: the EyeLink default saccade detection was used unchanged, and its thresholds are not tuned for microsaccade detection.

**4.6 Trial-level quality control.** A trial was retained if at least 80% of its samples carried a valid gaze position and no more than 20% of its valid samples fell outside a 3 deg window around fixation. Six of the 2,680 trials failed and 2,674 were retained, a loss of 0.22%. No trial failed the valid-sample criterion: the lowest valid fraction among the retained trials was 0.901, so the 80% threshold never bound. All six failures were failures of the fixation criterion, and they are listed individually in Table S3.3. They were contributed by three participants and fell in all five target classes, and in each of them between 20.6% and 45.5% of the valid samples lay outside the 3 deg window, against a median of 0.90% across retained trials. No participant lost more than three trials and no class lost more than two, and every one of the 35 participants contributed retained trials in all five classes.

**Table S3.3.** The six trials excluded by offline quality control, of 2,680 linked trials. The exclusion criterion was more than 20% of valid samples outside a 3 deg window around fixation; no trial fell below the 80% valid-sample threshold. Block index is zero-based, as in the archived files. No trial was excluded online, because no online rejection criterion was in force (Section 3.5).

| Participant | Block | Trial ID | Target class | Valid samples (%) | Valid samples outside 3 deg (%) |
|---|---|---|---|---|---|
| J010 | 1 | 20 | Letters | 97.7 | 28.2 |
| J022 | 0 | 6 | Control | 100.0 | 43.8 |
| J022 | 1 | 7 | Faces | 94.9 | 45.5 |
| J022 | 1 | 9 | Inverted face | 98.3 | 20.6 |
| J024 | 0 | 1 | Control | 99.4 | 21.8 |
| J024 | 0 | 3 | Glyph | 96.6 | 22.0 |

**Alt text:** The table lists the six trials excluded by offline gaze quality control, giving participant, block, trial ID, target class, percentage of valid samples, and percentage of valid samples outside the 3 deg fixation window.

**4.7 Statistical analysis.** All tests were computed on participant-level class means, so that trials were not treated as independent observations. The omnibus test of target class was a repeated-measures F statistic evaluated against 20,000 permutations of the class labels within each participant. Pairwise contrasts were two-sided paired sign-flip permutation tests, using 200,000 permutations for the eye and oculomotor measures and 2,000,000 permutations for the behavioural measures. The confidence interval for each paired mean difference was given by the 2.5th and 97.5th percentiles of 10,000 participant-resampled bootstrap draws. Holm's correction (Holm, 1979) was applied to three explicitly defined families: across the twelve eye/oculomotor omnibus tests of Table S3.5; within the three critical contrasts of each metric in Table S3.6; and, for the behavioural analyses, across all ten pairwise category contrasts within each metric. The random seed was 20260902. Analyses were run under Python 3.12.13 with pandas 2.2.3, NumPy 2.3.5 and Pillow 12.2.0.

### 5. Results

**5.1 Yield and data quality.** Across the 35 participants, the behavioural logs contained 2,740 trials from 137 blocks. EyeLink EDF files were available for 134 of those blocks, yielding 2,680 behaviourally linked eye-tracking trials; the remaining 60 behavioural trials came from three blocks for which no EDF file was written. Of these 2,680 linked trials, 2,674 passed quality control, a loss of 0.22%. The mean valid gaze fraction across retained trials was 0.988. The number of trials discarded because gaze left the fixation region is therefore zero online and six offline, 0.22% of the linked trials (Table S3.3). Nothing was rejected online because no online rejection criterion was in force. Fixation was enforced only by the gaze-contingent response gate of Section 3.5, which had to be satisfied before any response was recorded and which neither aborted nor flagged a trial thereafter, so that every trial contributes its full 30 s window whatever the gaze did after the gate opened. The window of interest used for the offline decision was 3 deg of visual angle around the screen centre, applied to the proportion of valid samples of a trial rather than to any single excursion.

**5.2 Fixation stability and gaze position.** Table S3.4 gives participant-level means for all five target classes and Table S3.5 the omnibus tests. Fixation was tight in every class: roughly 99% of samples carried a valid gaze position, mean gaze distance from fixation was between 0.81 and 0.83 deg, and between 1.30% and 1.41% of valid samples fell outside the 3 deg window. Target class had no reliable effect on the proportion of samples outside the 1.5, 2 or 3 deg windows, on mean gaze distance from fixation, or on the horizontal or vertical spread of gaze position (Figures S3.3 to S3.5).

One qualification is required. The omnibus test of valid gaze fraction returned $F(4, 136) = 2.44$, $p = .048$ before correction and $p = .574$ after Holm correction across the twelve omnibus tests, and the corresponding faces-versus-control contrast was $p = .046$ before and $p = .139$ after correction within its family of three. The difference is 0.00099, one tenth of one percentage point of valid samples. Valid gaze fraction is a measure of track loss rather than of oculomotor behaviour, and the direction of the difference, marginally more usable data on face trials, is not one that could manufacture more frequent disappearances. We report it because it is the only omnibus test in Table S3.5 that falls below .05 uncorrected, and it should not be read as evidence of a category effect.

**5.3 Blinks and saccades.** EyeLink EBLINK event-record rate lay between 0.232 and 0.247 records per second and ESACC event-record rate between 1.97 and 2.01 records per second, with mean saccade amplitudes between 0.74 and 0.80 deg. Neither event-record rate showed a reliable effect of target class, and neither did saccade amplitude (Table S3.5, Figures S3.6 and S3.7). Within the amplitude family the largest contrast is faces against letters, −0.034 deg, $p = .054$ uncorrected and $p = .161$ after Holm correction within the three contrasts of that measure; it does not involve the control class and we draw no conclusion from it. The paired contrasts in Table S3.6 provide interval estimates for any residual difference: for EBLINK event-record rate the 95% bootstrap interval for the faces-minus-control difference runs from −0.033 to +0.003 records per second, and for ESACC event-record rate from −0.094 to +0.092 records per second, against class means of roughly 0.24 and 1.99 respectively.

**5.4 Pupil size.** Pupil measures are reported as exploratory. Neither mean pupil area nor baseline-corrected pupil area showed a reliable effect of target class (Table S3.5, Figure S3.8), and none of the critical contrasts approached significance (Table S3.6). No pupil hypothesis was specified in advance, and baseline-corrected pupil area depends on the choice of baseline interval; these results are therefore reported descriptively, and no inference is drawn from them.

**5.5 Behavioural measures in the control dataset.** Table S3.7 and Figures S3.9 and S3.10 give the behavioural measures for completeness, since they were computed from the same retained trials. Target class had no reliable effect on any of them, as already stated in Section 2.1. Two of the ten pairwise contrasts on total invisible duration reach $p < .05$ uncorrected, control versus faces at $p = .014$ and control versus letters at $p = .006$, but neither survives Holm correction across the ten contrasts of that measure (adjusted p = .123 and .060), and the omnibus test does not reach significance. For the rate and invisibility-fraction summaries, the denominator is the response-enabled part of the 30 s trial rather than the full stimulus window, matching the fact that responses could be recorded only after the fixation gate had opened. Summed invisible duration still excludes any disappearance that was ongoing at the end of the trial, whereas the invisibility fraction of the main text includes such right-censored intervals at their observed length.

**Table S3.4.** Participant-level means (SEM) by target class, computed over the 2,674 retained trials of 35 participants. Percentages are of valid gaze samples within the 30 s analysis window. Pupil area is in EyeLink arbitrary units.

| Measure | Control | Faces | Letters | Glyphs | Inverts |
|---|---|---|---|---|---|
| Valid gaze fraction | 0.9883 (0.0017) | 0.9893 (0.0017) | 0.9888 (0.0018) | 0.9889 (0.0016) | 0.9880 (0.0019) |
| Outside 1.5 deg (%) | 7.522 (1.606) | 7.247 (1.705) | 8.451 (2.200) | 7.392 (1.791) | 7.222 (1.737) |
| Outside 2 deg (%) | 2.614 (0.362) | 2.986 (0.700) | 3.334 (0.813) | 3.076 (0.700) | 2.825 (0.637) |
| Outside 3 deg (%) | 1.409 (0.162) | 1.304 (0.175) | 1.411 (0.165) | 1.405 (0.163) | 1.336 (0.154) |
| Mean gaze distance (deg) | 0.816 (0.048) | 0.813 (0.047) | 0.830 (0.052) | 0.810 (0.048) | 0.816 (0.049) |
| SD gaze x (deg) | 0.387 (0.023) | 0.383 (0.024) | 0.386 (0.023) | 0.387 (0.023) | 0.387 (0.024) |
| SD gaze y (deg) | 0.894 (0.075) | 0.864 (0.078) | 0.887 (0.076) | 0.907 (0.072) | 0.899 (0.077) |
| EyeLink EBLINK event-record rate (records/s) | 0.247 (0.033) | 0.232 (0.033) | 0.243 (0.034) | 0.243 (0.030) | 0.244 (0.034) |
| EyeLink ESACC event-record rate (records/s) | 1.987 (0.155) | 1.982 (0.160) | 2.002 (0.166) | 2.006 (0.156) | 1.974 (0.153) |
| Mean saccade amplitude (deg) | 0.766 (0.039) | 0.740 (0.041) | 0.774 (0.043) | 0.800 (0.048) | 0.747 (0.035) |
| Mean pupil area | 1315.8 (57.5) | 1324.8 (56.9) | 1313.1 (53.4) | 1340.7 (56.7) | 1326.6 (57.6) |
| Baseline-corrected pupil | 36.0 (15.5) | 23.2 (16.2) | 17.9 (16.7) | 25.6 (18.3) | 32.1 (15.4) |

**Alt text:** The table gives participant-level means with SEMs for Control, Faces, Letters, Glyphs, and Inverts across gaze-validity, fixation, blink, saccade, and pupil measures.

**Table S3.5.** Omnibus tests of target class across all five classes, computed on participant-level means with 20,000 within-participant label permutations. The final column applies Holm's correction across the twelve tests.

| Measure | F(4, 136) | p | p (Holm, 12) |
|---|---|---|---|
| Valid gaze fraction | 2.44 | .048 | .574 |
| Outside 1.5 deg (%) | 1.07 | .382 | 1.000 |
| Outside 2 deg (%) | 1.18 | .341 | 1.000 |
| Outside 3 deg (%) | 0.87 | .493 | 1.000 |
| Mean gaze distance (deg) | 0.73 | .581 | 1.000 |
| SD gaze x (deg) | 0.07 | .990 | 1.000 |

| Measure | F(4, 136) | p | p (Holm, 12) |
|---|---|---|---|
| SD gaze y (deg) | 1.75 | .138 | 1.000 |
| EyeLink EBLINK event-record rate (records/s) | 0.79 | .539 | 1.000 |
| EyeLink ESACC event-record rate (records/s) | 0.21 | .937 | 1.000 |
| Mean saccade amplitude (deg) | 2.19 | .066 | .725 |
| Mean pupil area | 2.05 | .085 | .853 |
| Baseline-corrected pupil | 0.71 | .595 | 1.000 |

**Alt text:** The table lists omnibus repeated-measures test results for twelve gaze, oculomotor, and pupil measures across all five target classes, with F statistics, uncorrected p values, and Holm-adjusted p values.

**Table S3.6.** Paired contrasts among the three critical target classes for the six gaze, blink and saccade measures plotted in Figure S3.2 and for baseline-corrected pupil area, which is exploratory (Section 5.4). Differences are participant-level paired means, intervals are 95% bootstrap intervals over 10,000 participant resamples, p values are from two-sided sign-flip permutation tests with 200,000 permutations, and the Holm correction is applied within the three contrasts of each measure.

| Measure | Contrast | Difference | 95% CI | p | p (Holm) |
|---|---|---|---|---|---|
| Valid gaze fraction | Faces − control | +0.00099 | [+0.00008, +0.00196] | .046 | .139 |
| | Letters − control | +0.00045 | [−0.00058, +0.00150] | .405 | .405 |
| | Faces − letters | +0.00054 | [−0.00023, +0.00135] | .197 | .395 |
| Outside 3 deg (%) | Faces − control | −0.106 | [−0.252, +0.057] | .199 | .397 |
| | Letters − control | +0.002 | [−0.123, +0.135] | .976 | .976 |
| | Faces − letters | −0.108 | [−0.194, −0.018] | .025 | .076 |
| Mean gaze distance (deg) | Faces − control | −0.003 | [−0.034, +0.026] | .839 | .839 |
| | Letters − control | +0.015 | [−0.017, +0.048] | .396 | .792 |
| | Faces − letters | −0.018 | [−0.045, +0.008] | .197 | .590 |
| EyeLink EBLINK event-record rate (records/s) | Faces − control | −0.015 | [−0.033, +0.003] | .108 | .324 |
| | Letters − control | −0.004 | [−0.022, +0.012] | .625 | .625 |
| | Faces − letters | −0.011 | [−0.028, +0.006] | .234 | .468 |
| EyeLink ESACC event-record rate (records/s) | Faces − control | −0.005 | [−0.094, +0.092] | .919 | 1.000 |
| | Letters − control | +0.015 | [−0.067, +0.092] | .723 | 1.000 |
| | Faces − letters | −0.020 | [−0.110, +0.061] | .659 | 1.000 |
| Mean saccade amplitude (deg) | Faces − control | −0.026 | [−0.066, +0.012] | .218 | .435 |
| | Letters − control | +0.008 | [−0.036, +0.056] | .764 | .764 |
| | Faces − letters | −0.034 | [−0.068, −0.002] | .054 | .161 |
| Baseline-corrected pupil | Faces − control | −12.82 | [−41.72, +11.99] | .385 | .770 |
| | Letters − control | −18.12 | [−44.14, +6.22] | .175 | .526 |

| Measure | Contrast | Difference | 95% CI | p | p (Holm) |
|---|---|---|---|---|---|
| | Faces − letters | +5.30 | [−18.29, +28.94] | .664 | .770 |

**Alt text:** The table lists paired contrasts among Faces, Letters, and Control for gaze, blink, saccade, and baseline-corrected pupil measures. For each contrast it gives the paired mean difference, 95% bootstrap confidence interval, uncorrected p value, and within-measure Holm-adjusted p value.

**Table S3.7.** Behavioural measures of the control dataset, participant-level means over the same retained trials, with the omnibus test across all five classes.

| Measure | Control | Faces | Letters | Glyphs | Inverts | p |
|---|---|---|---|---|---|---|
| Invisibility fraction (response-enabled) | 0.113 | 0.103 | 0.103 | 0.109 | 0.106 | .105 |
| Disappearance onsets (/s response-enabled visible) | 0.143 | 0.136 | 0.137 | 0.141 | 0.138 | .687 |
| Disappearance reports (/trial) | 3.32 | 3.19 | 3.24 | 3.28 | 3.23 | .812 |
| Reappearance reports (/trial) | 3.07 | 2.90 | 3.01 | 3.02 | 2.94 | .510 |
| Total invisible duration (s/trial) | 3.16 | 2.88 | 2.90 | 3.07 | 2.96 | .078 |

**Alt text:** The table gives participant-level behavioural means for Control, Faces, Letters, Glyphs, and Inverts, with an omnibus p value for each measure. Measures include response-enabled invisibility fraction, disappearance onset rate, disappearance and reappearance reports per trial, and total invisible duration.

### 5.6 Graphical summaries

Figure S3.2 collects the category contrasts of Sections 5.2 and 5.3 in a single panel. Each point is the paired participant-level difference between a target class and the control class, divided by the between-participant standard deviation of the control class, so that measures recorded in different units can be read against one axis; the interval is the 95% bootstrap interval of Table S3.6 expressed in the same units. No estimate in the figure exceeds 0.12 control standard deviations, and nine of the twelve intervals lie entirely within ±0.2. The largest absolute interval endpoint is 0.29, for mean saccade amplitude on face trials, where the estimate is negative, that is, in the direction of smaller rather than larger eye movements for faces. The pupil measures of Section 5.4 are not plotted, because they are exploratory and are not a mechanism by which a target category could displace the retinal image; they are reported in Tables S3.4 to S3.6.

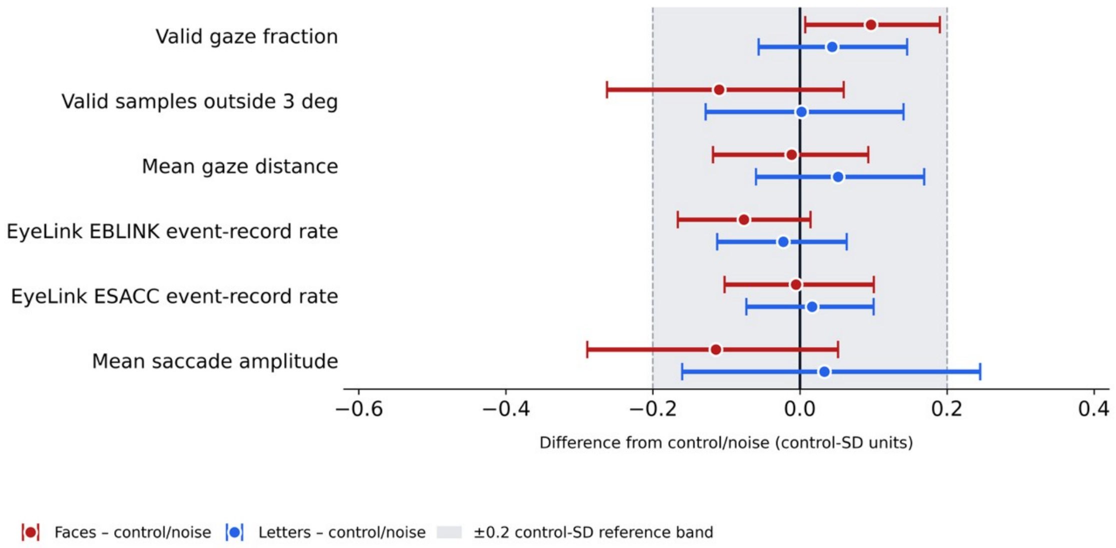


**Figure S3.2.** *Standardized category contrasts across the eye measures.* Paired differences from the control class in units of the between-participant standard deviation of the control class, with 95% percentile bootstrap intervals over 10,000 participant resamples. The shaded band marks ±0.2 control standard deviations. Pupil measures are exploratory and are not plotted; they are given in Tables S3.4 to S3.6 and Section 5.4. The underlying differences, intervals and p values are those of Table S3.6. Positive values indicate a larger value for the target class than for control.

**Alt text:** A horizontal interval plot has six rows for valid gaze fraction, samples outside 3 deg, mean gaze distance, EyeLink EBLINK event-record rate, EyeLink ESACC event-record rate, and mean saccade amplitude. Red points and horizontal intervals represent Faces minus control/noise, blue points represent Letters minus control/noise, a vertical line marks zero, and a grey vertical band marks ±0.2 control-SD units.

The remaining figures show the participant-level means that underlie Tables S3.4 and S3.7. In every panel the open symbols are the 35 participant means, jittered horizontally for visibility, the coloured bar is the group mean and the black error bars give ±1 SEM. All five target classes are plotted. Where the measure has a meaningful zero the axis is anchored at it, so that the size of the differences between classes is not visually magnified.

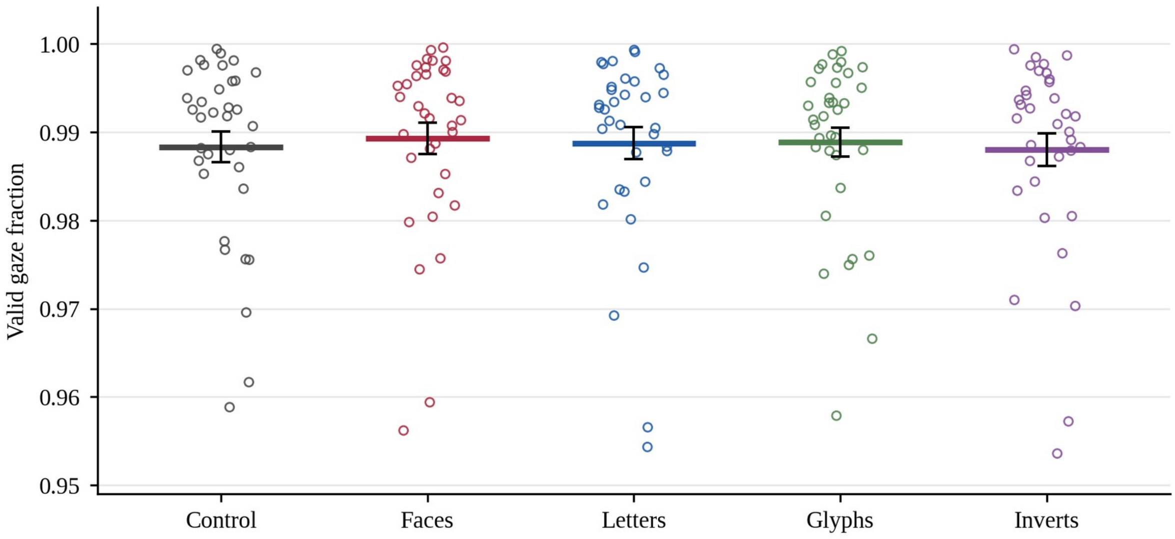


**Figure S3.3.** *Valid gaze fraction by target class.* The proportion of samples in the 30 s analysis window carrying a finite gaze position in at least one eye.

**Alt text:** A jittered open-circle scatter plot shows valid gaze fraction for Control, Faces, Letters, Glyphs, and Inverts. Each category has many participant points, a coloured horizontal mean bar, and black vertical error bars; values are concentrated near the top of the axis around 0.99.

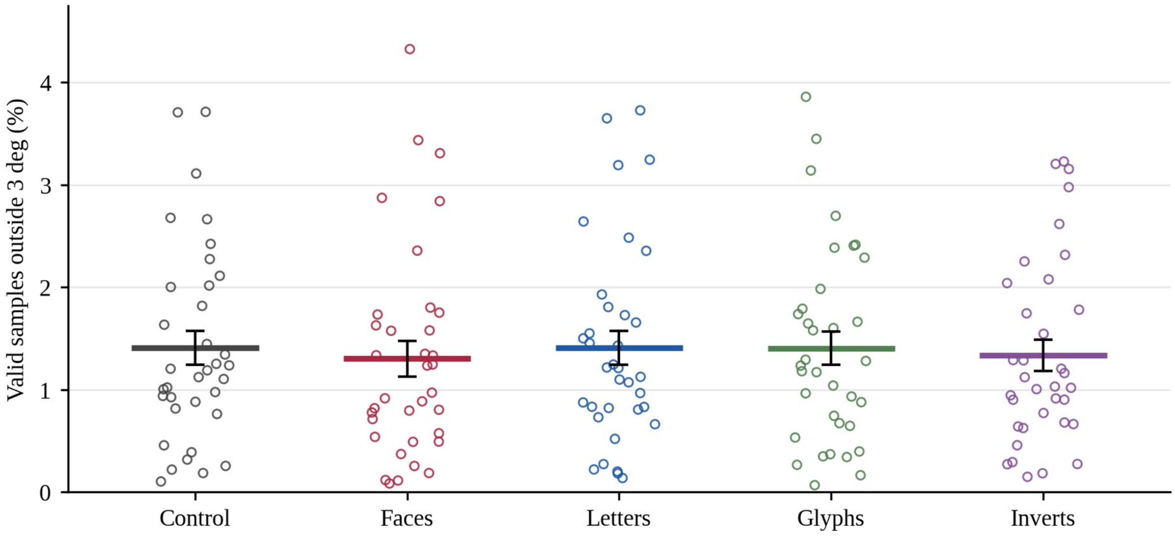


**Figure S3.4.** *Valid gaze samples outside the 3 deg fixation window.* The percentage of valid samples farther than 3 deg from the screen centre, the criterion used for trial-level quality control.

**Alt text:** A jittered open-circle scatter plot shows the percentage of valid samples outside the 3 deg fixation window for Control, Faces, Letters, Glyphs, and Inverts. Each category has participant points, a coloured mean bar, and black error bars, with most points low on the axis and a smaller number extending upward.

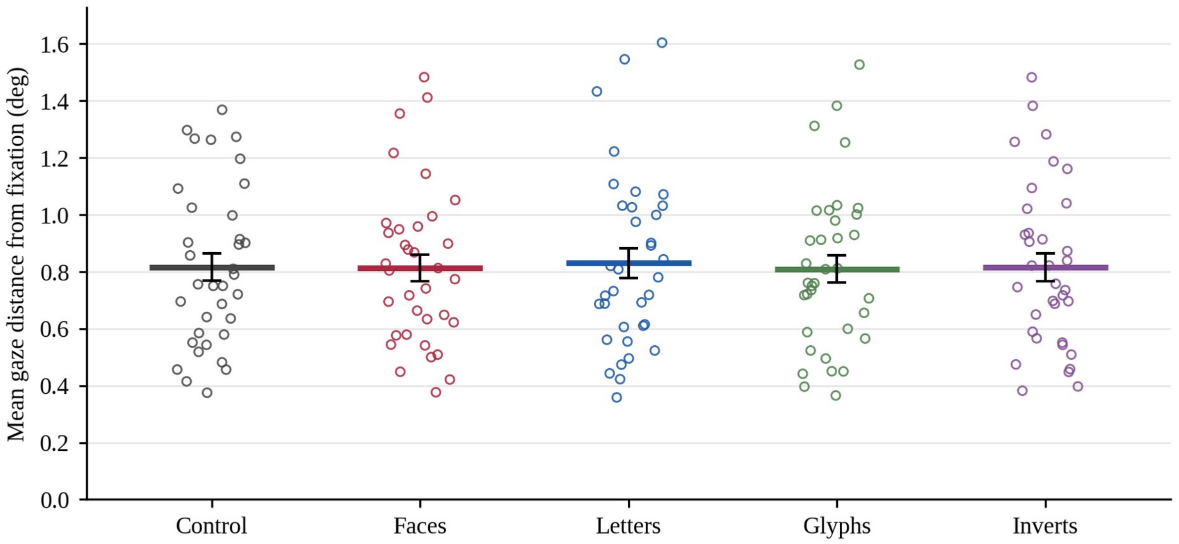


**Figure S3.5.** *Mean gaze distance from fixation.* The mean distance of valid gaze samples from the screen centre, in degrees of visual angle.

**Alt text:** A jittered open-circle scatter plot shows mean gaze distance from fixation, in degrees, across Control, Faces, Letters, Glyphs, and Inverts. Coloured horizontal bars mark category means, black vertical bars show errors, and participant points are spread vertically around the mean of each category.

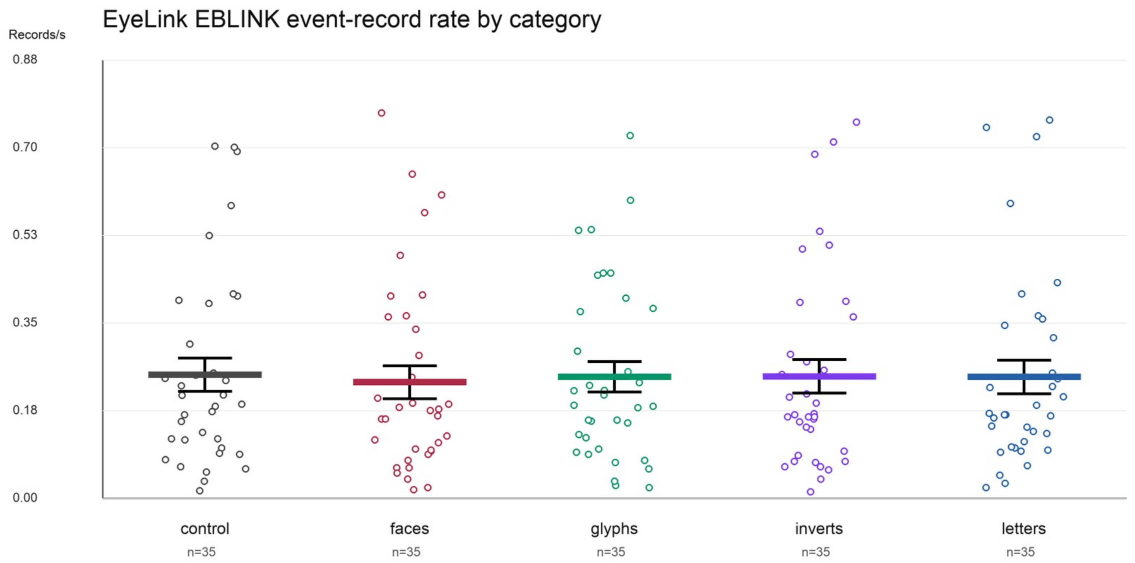


**Figure S3.6.** *EyeLink EBLINK event-record rate by target class. EBLINK records overlapping the analysis window, expressed as records per second.*

**Alt text:** A jittered open-circle scatter plot shows EyeLink EBLINK event-record rate, in records per second, across Control, Faces, Glyphs, Inverts, and Letters. Each category has a coloured horizontal mean bar and black error bars, with participant points spread from low rates to a smaller number of higher values.

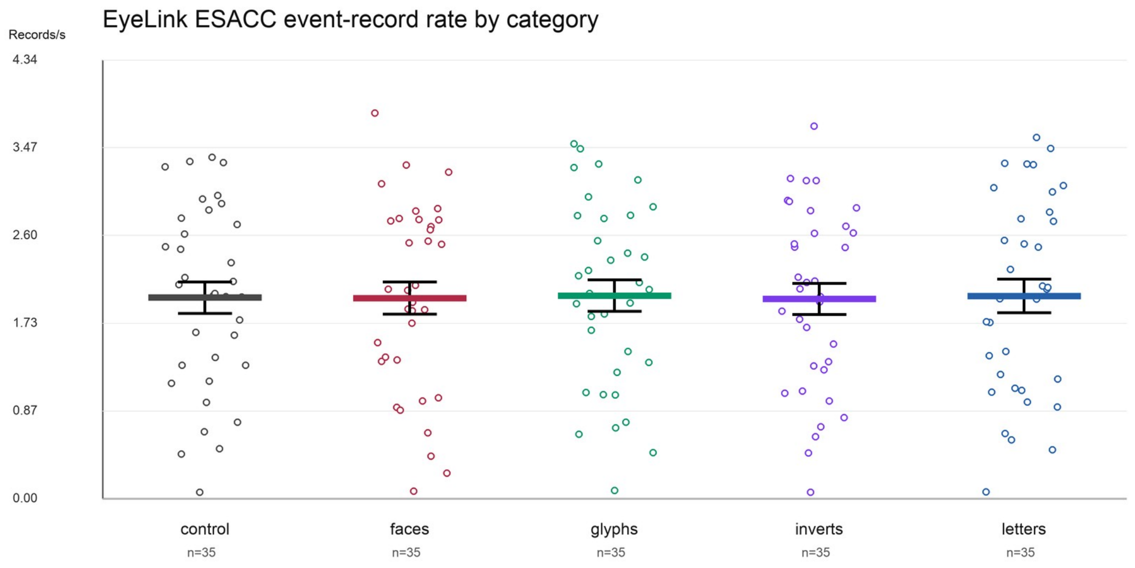


**Figure S3.7.** *EyeLink ESACC event-record rate by target class. ESACC records overlapping the analysis window, expressed as records per second.*

**Alt text:** A jittered open-circle scatter plot shows EyeLink ESACC event-record rate, in records per second, across Control, Faces, Glyphs, Inverts, and Letters. Participant points are spread broadly over the vertical axis, with coloured horizontal mean bars and black vertical error bars for each category.

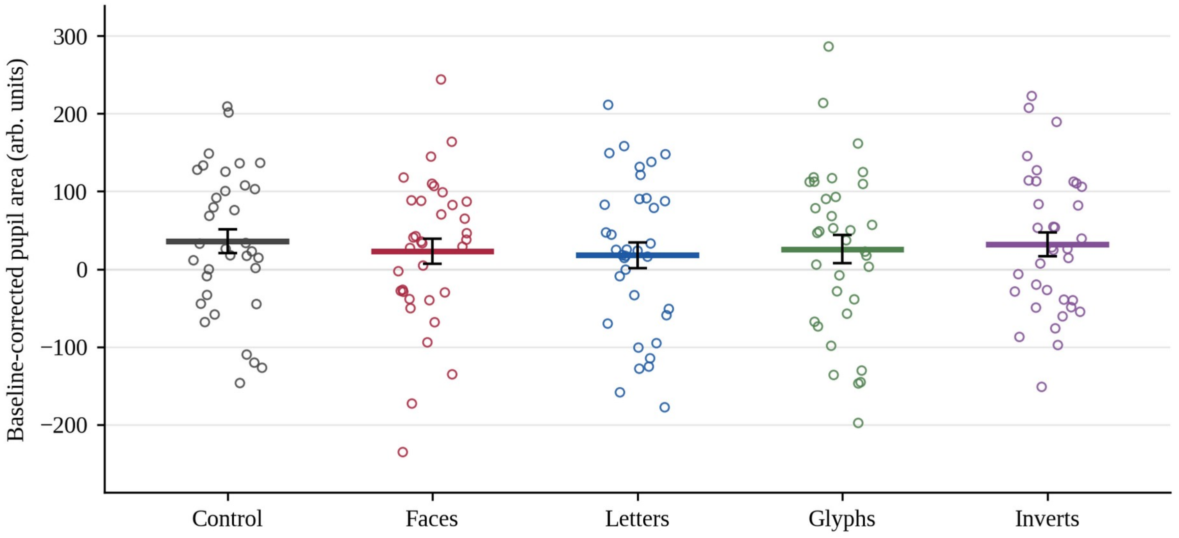


**Figure S3.8.** *Baseline-corrected pupil area by target class.* The mean pupil area over the analysis window minus the mean over its first 500 ms, in EyeLink arbitrary units. The grey line marks no change from baseline. These measures are exploratory.

**Alt text:** A jittered open-circle scatter plot shows baseline-corrected pupil area for Control, Faces, Letters, Glyphs, and Inverts. Participant points extend above and below a horizontal grey zero line; coloured horizontal bars show category means and black vertical bars show errors.

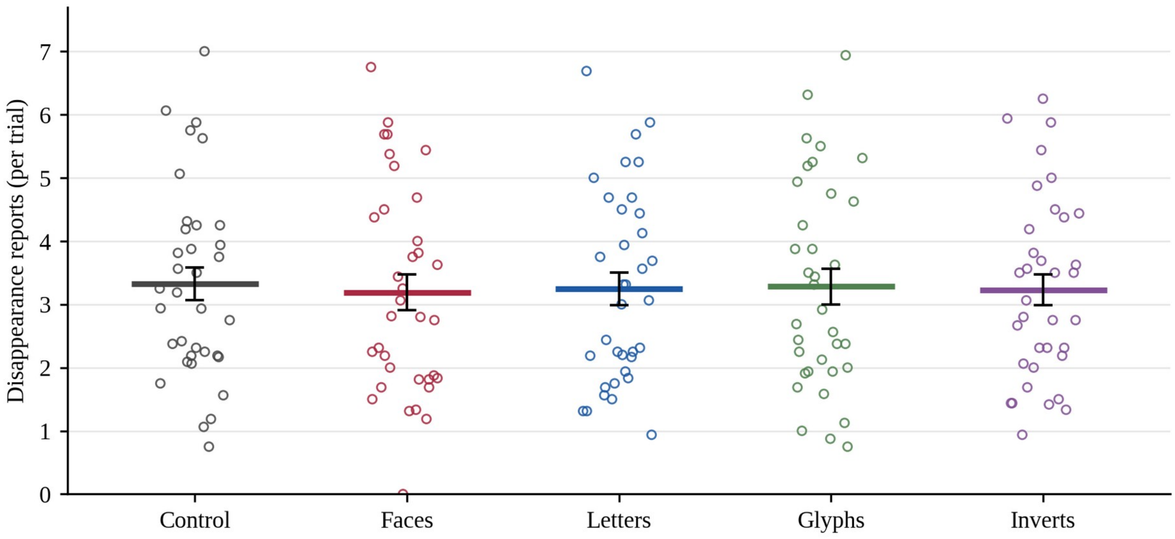


**Figure S3.9.** *Disappearance reports by target class.* Reported disappearances per trial in the control dataset.

**Alt text:** A jittered open-circle scatter plot shows disappearance reports per trial for Control, Faces, Letters, Glyphs, and Inverts. Participant points range from near zero to several reports per trial, with coloured horizontal mean bars and black vertical error bars for each category.

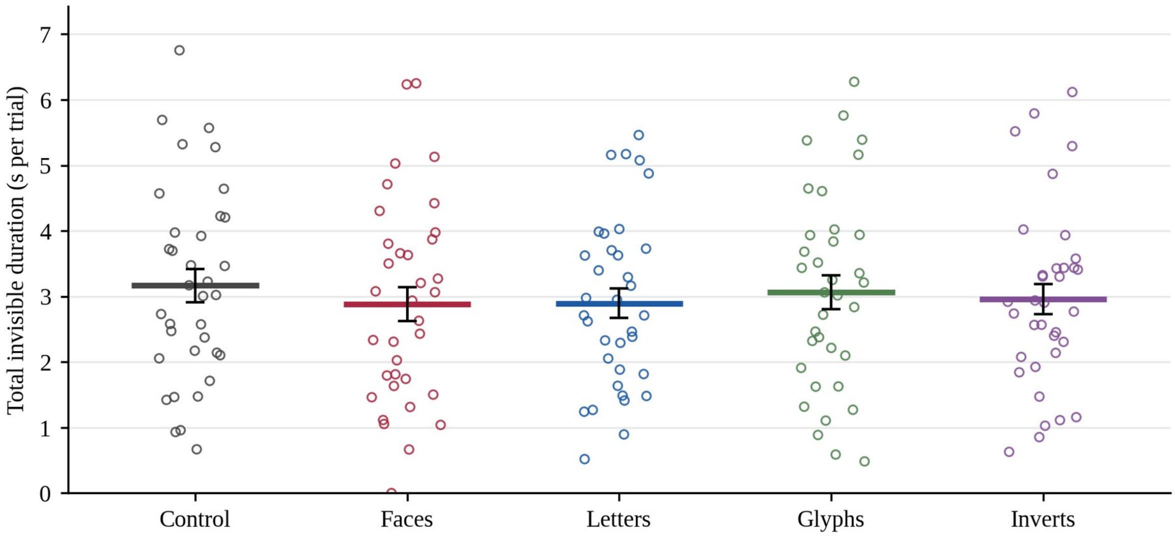


**Figure S3.10.** *Total invisible duration by target class.* The summed duration of reported invisible episodes per trial in the control dataset. Episodes still ongoing at the end of a trial are not included.

**Alt text:** A jittered open-circle scatter plot shows total invisible duration per trial for Control, Faces, Letters, Glyphs, and Inverts. Participant points range from near zero to longer durations, with coloured horizontal mean bars and black vertical error bars for each category.

### 6. What the dataset supports, and what it does not

The dataset supports one limited conclusion. In an independent MIB experiment run with the same eye tracker, at the same target eccentricity, and within an inducer field of nearly the same size, gaze position, fixation stability, EyeLink EBLINK event-record rate, EyeLink ESACC event-record rate and saccade amplitude showed no reliable differences between phase-randomized control, face and letter targets (Figure S3.2); the pupil measures, which are exploratory, likewise showed none. The simplest oculomotor account of category differences in MIB dynamics (one in which faces or letters draw the eye differently, are fixated less steadily, or are accompanied by systematically different EBLINK or ESACC event-record rates) therefore has no support in the only comparable gaze record available to us.

Four limits bound that claim.

First, these are not Experiment B's eye movements. The argument is an argument from a closely related design, not from the same observers or the same trials.

Second, all measures reported here are tonic. They summarize the whole 30 s window and cannot address transition-locked effects, which is precisely where the literature locates the relationship between fixational eye movements and MIB (Bonneh et al., 2010; Hsieh & Tse, 2009). A transition-locked analysis was not attempted. It was not planned for this dataset and no event-alignment procedure was validated for it, and the perceptual reports are subjective and coarse in time relative to the fixational events of interest. Response times were additionally logged relative to the fixation gate rather than to image onset; Section 4.3 shows that the two time bases can be reconciled, so this is a practical complication rather than the reason the analysis was not attempted.

Third, the behavioural class effect was itself absent in this dataset (Section 2.1). The analysis is therefore a control on the stimulus classes under the conditions of this dataset rather than on the behavioural effect: it shows that these three classes, at this eccentricity and within this inducer, produced no reliable differences on the oculomotor measures recorded here. It does not establish that the class difference observed in Experiment B is unaccompanied by an oculomotor difference. What it addresses is the version of the confound in which the classes are looked at differently irrespective of what the observer reports: on these measures and under these conditions no reliable category difference was detected, and the endpoints of the 95% bootstrap intervals in Figure S3.2 lie within approximately ±0.29 control standard deviations.

Fourth, microsaccades were not extracted, the pupil analyses are exploratory for the reasons given in Section 5.4 and are not plotted in Figure S3.2, and the widest interval in that figure remains compatible with a difference of approximately 0.29 control standard deviations in mean saccade amplitude.

### 7. Archived files

The stimulus set, the processed data, the results, and the analysis and figure code of this control experiment are archived with this supplement in the folder MIB_eye_tracking_control_experiment_analysis of the study repository. The raw EyeLink EDF recordings are available from the authors on reasonable request.

### 8. References

All works cited in this supplement are listed in the consolidated reference list of the main manuscript.
